# The neurotrophin DNT-2 regulates cell survival and connectivity via the Toll-2 receptor during visual system development of *Drosophila*

**DOI:** 10.1101/2025.07.18.665476

**Authors:** Naser Alshamsi, Francisca Rojo-Cortés, Bangfu Zhu, Samaher Fahy, Guiyi Li, Anna Lassota, Marta Moreira, Alicia Hidalgo

## Abstract

During development, neurons are produced in excess and those that receive trophic support are maintained, whereas excess neurons are eliminated, enabling the establishment of appropriate neural circuits. In vertebrates, neurotrophin ligands promote cell survival during periods of naturally occurring cell death, by signalling through p75 and Trk receptors. In the *Drosophila* optic lobe, a wave of apoptosis occurs during neural circuit development; however, whether this also involves neurotrophism remains unresolved. *Drosophila* neurotrophins (DNTs) are encoded by *spätzle* (spz) paralogue genes and bind Toll receptors instead. Here, we focused on DNT-3 (previously known as spz-3) and *DNT-2* (also known as *spz-5*) to ask whether they underlie neurotrophism in visual system development. We show that *DNT-3* (*spz-3*) and *DNT-2* (*spz-5*) are both expressed in the retina and in medulla neurons, and multiple Tolls are expressed across lamina and medulla neurons. Over-expression of *DNT-3* (*spz-3*) and *DNT-2* (*spz-5*) could rescue natural occurring cell death, whereas their loss of function caused cell death, showing that DNT-3 and *DNT-2* can, and are required to, promote cell survival during optic lobe development. Importantly, *DNT-2* is expressed in Mi1 neurons and *Toll-2* in connecting L1 neurons. We show that DNT-2 functions in concert with Toll-2, as *Toll-2* RNAi knock-down prevented the rescue of apoptosis by *DNT-2* over-expression and all Toll-2+ neurons were lost in *DNT-2* mutants. Furthermore, alterations in *DNT-2* or *Toll-2* expression levels impaired connectivity of L1 neurons at the M1 medulla layer and altered dendritic morphology of L1 neurons. These data suggest that L1 neurons could take up *DNT-2* secreted from medulla neurons during the establishment of connectivity patterns. As *DNT-3* (*spz-3*) and *DNT-2* (*spz-5*) are expressed in the medulla and they could influence both lamina and medulla neurons, this suggests that their function maintaining cell survival could enable the stabilisation or alignment of connected neurons across medulla columns.

## Introduction

Neural circuits emerge during growth, presenting the challenge of how cell number is controlled to ensure circuits drive appropriate brain function and behaviour. During development, neurons and glia are generated in excess, and excess cells are eliminated by apoptosis enabling appropriate connectivity and neural circuit formation (Levi-Montalcini, 1987, Raff et al., 1993). Neuronal survival is maintained by neurotrophic factors secreted in limited amounts by target cells, leading to the survival of only those neurons that receive trophic support (Levi-Montalcini, 1987, Davies, 2003). Similarly, glial cell survival is maintained by gliatrophic factors released by neurons (Raff et al., 1993). Neurons and glia that fail to receive trophic support and establish appropriate connectivity are eliminated by apoptosis, and in this way, cell number in interacting cell populations is adjusted in development (Levi-Montalcini, 1987, Raff et al., 1993). In this context, if neurotrophism is fundamental for nervous system development, it could have been enabled by evolutionarily conserved molecular mechanisms.

Neurotrophins – NGF, BDNF, NT3, NT4 - are the main growth factors maintaining neuronal survival in the vertebrate nervous system (Levi-Montalcini, 1987, Lu et al., 2005). Importantly, they can also promote cell death, depending on context (Lu et al., 2005). They can promote cell survival via their Trk receptors and ERK and AKT downstream and via p75^NTR^ and NFκB downstream, or cell death via p75^NTR^, Sortilin, and JNK signalling instead (Lu et al., 2005). Homologues of neurotrophins and their receptors have been found across the invertebrates (Foldi et al., 2017, McIlroy et al., 2013, Zhu et al., 2008, DeLotto and DeLotto, 1998, Benito-Gutierrez et al., 2005, Laramore et al., 2011, Lauri et al., 2016, Hallbook, 1999), but functional in vivo evidence for their involvement in the regulation of cell survival or cell death outside the vertebrates remains limited. In the fruit-fly, *Drosophila* neurotrophins (DNTs) are encoded by *spätzle* (*spz*) paralogue genes, that include sequence, structural and functional neurotrophin homologues (DeLotto and DeLotto, 1998, Foldi et al., 2017, Zhu et al., 2008). DNT ligands bind the Trk-family Kekkon (Kek) receptors, and also Toll receptors (Foldi et al., 2017, McIlroy et al., 2013, Ulian-Benitez et al., 2017, Weber et al., 2003). Spz-1 is the well-known ligand of Toll-1 (also known as Toll), DNT-1 (also known as Spz-2) and DNT-2 (also known as Spz-5) of Toll-7 and Toll-6 respectively, and DNT-3 (also known as Spz-3) is a candidate ligand for Toll-8 (Ballard et al., 2014, McIlroy et al., 2013, Weber et al., 2003). DNT-1 and -2 and their Toll-6 and -7 receptors can promote neuronal survival in embryonic, pupal and larval ventral nerve cords and in adult brains, demonstrating functional evolutionary conservation (Foldi et al., 2017, McIlroy et al., 2013, Sun et al., 2024, Zhu et al., 2008). There are six *spz* and nine *Toll* paralogous genes in Drosophila, which could play distinct functions. In fact, at least full-length DNT-1 and Toll-1 can promote cell death instead, and at least Toll-6 can promote either cell survival or cell death, depending on context (Foldi et al., 2017, Singh et al., 2025, Zhu et al., 2008) . Importantly, mature DNT-1 and DNT-2 with Toll-6 and Toll-7 are required for and can promote neuronal survival during circuit formation in the embryonic ventral nerve cord (McIlroy et al., 2013, Zhu et al., 2008).

The developing visual system is an ideal context in which to further test neurotrophism and its link to neural circuit formation in Drosophila. The optic lobes of adult flies are formed from neurogenesis during larval development, followed by a wave of cell death taking place in the pupa, at the time of neural circuit formation (Holguera and Desplan, 2018, Togane et al., 2012). Apoptosis is partly induced by ecdysone at the onset of metamorphosis, driving clearance of larval cells that do not contribute to the adult brain, but there is also a significant extent of ecdysone-independent cell death (Hara et al., 2013). Apoptosis initiates just after pupariation, reaching a peak at 24h after puparium formation (APF), and decreasing by 48h APF (Togane et al., 2012, Hara et al., 2013). During this time (24-50h APF), connectivity between photoreceptors, lamina and medulla neurons is established; this is followed by medulla neurons connecting to lobula neurons; and by 72h APF cell death has greatly diminished and synaptogenesis completes connectivity patterns, in preparation for adult eclosion at 96h APF (Millard and Pecot, 2018, Melnattur and Lee, 2011, Hadjieconomou et al., 2011, Kurmangaliyev et al., 2020). Thus, 24-48h APF is a critical period to maintain necessary lamina and medulla neurons alive in the optic lobe.

The development of the *Drosophila* visual system has been well described (Holguera and Desplan, 2018, Melnattur and Lee, 2011, Millard and Pecot, 2018, Hadjieconomou et al., 2011, Behnia and Desplan, 2015). The retina of the compound eye consists of 750-800 ommatidia each housing eight photoreceptor cells (R1-R8), that form precise connections with target neurons in the optic lobes. R1-R6 target to the lamina, making synaptic contact with L1-L3 lamina neuron dendrites, and together with L4 and L5, organize into lamina cartridges (Tuthill et al., 2014). R7 and R8, together with lamina neurons target to medulla layers M6 and M3, respectively, organizing into medullar columns that respond to the same point in visual space and maintain retinotopy. Medullar interneurons also form connections across multiple layers, where each layer represents different visual features (Fischbach and Hiesinger, 2008, Millard and Pecot, 2018). Medulla neurons connect with the lamina for feedback and with lobula and lobula plate neurons for higher-order visual processing, including motion detection, spatial orientation, and feature extraction (Behnia and Desplan, 2015, Borst et al., 2020, Courgeon and Desplan, 2019b). Neurons within the lobula complex integrate signals from the medulla and project to the optic glomeruli in the central brain and motor outputs to enable appropriate behavior (Behnia and Desplan, 2015, Borst et al., 2020, Courgeon and Desplan, 2019b). The temporal correlation between the pupal wave of cell death and neural circuit formation prompted the question of whether neurotrophism could be involved in visual system development.

Here, we asked whether neurotrophin family ligands encoded by *DNTs* (*spzs*) and their Toll receptors could regulate cell survival during neural circuit formation, in the *Drosophila* pupal optic lobe. Spz-5 is well known as Drosophila neurotrophin-2 (DNT-2), and as Spz-3 has been proposed to have neurotrophin functions which we expand on and demonstrate here, we refer to Spz-3 as DNT-3 (Zhu et al., 2008, Coutinho-Budd et al., 2017, Sun et al., 2024, Ballard et al., 2014, Ulian-Benitez et al., 2017).

## Results

### Differential expression of spz paralogues and Toll receptors in the developing optic lobe

To ask whether *DNTs* (s*pzs*) are expressed in the pupal optic lobe, we generated *T2A-Gal4* driver fly lines, crossed them to *10xUASmyrGFP* or *20UAS6xmCherry* reporter flies, and analysed resulting progeny optic lobes with anti-GFP antibodies, as required (Figure 1A). *spz-1*^*MIO2318*^*-T2A>myrGFP* and *spz-1*^*MIO2318*^*-T2A>6xm*Cherry revealed expression in a few centrifugal neurons in the lobula complex that projected to the lamina, subsequently medulla neurons and abundant arborisations into the lobula complex and medulla. Expression from *spz4*^*MI5678*^*-T2A->myrGFP* was not detected until 72h APF and then was found in the medulla and lobula complex and followed by the trachea. *spz-3-T2A>6xm*Cherry (hereby named DNT-3) was highly expressed in non-neuronal retinal cells and medulla neurons; and subsequently in the trachea and possibly glia. Finally, *DNT-2-T2A>6xmCherry (spz-5)* was found in medulla neurons, which could be tentatively identified as Mi1 medulla neurons (Nern et al., 2025) by 48h APF, and this pattern was maintained. Abundant cells expressed *spz-1* in the lobula complex (Figure 1B, left) and medulla (Figure 1B, right), and *DNT-3 (spz-3)* and *DNT-2 (spz-5)* in the medulla (Figure 1B) during optic lobe development. *DNT-3 (spz-3)* and *DNT-2 (spz-5)* were expressed in distinct non-neuronal cells in the retina, seen in Multi Colour Flip Out (MCFO) clones (Figure 1C).

**Figure 1.**
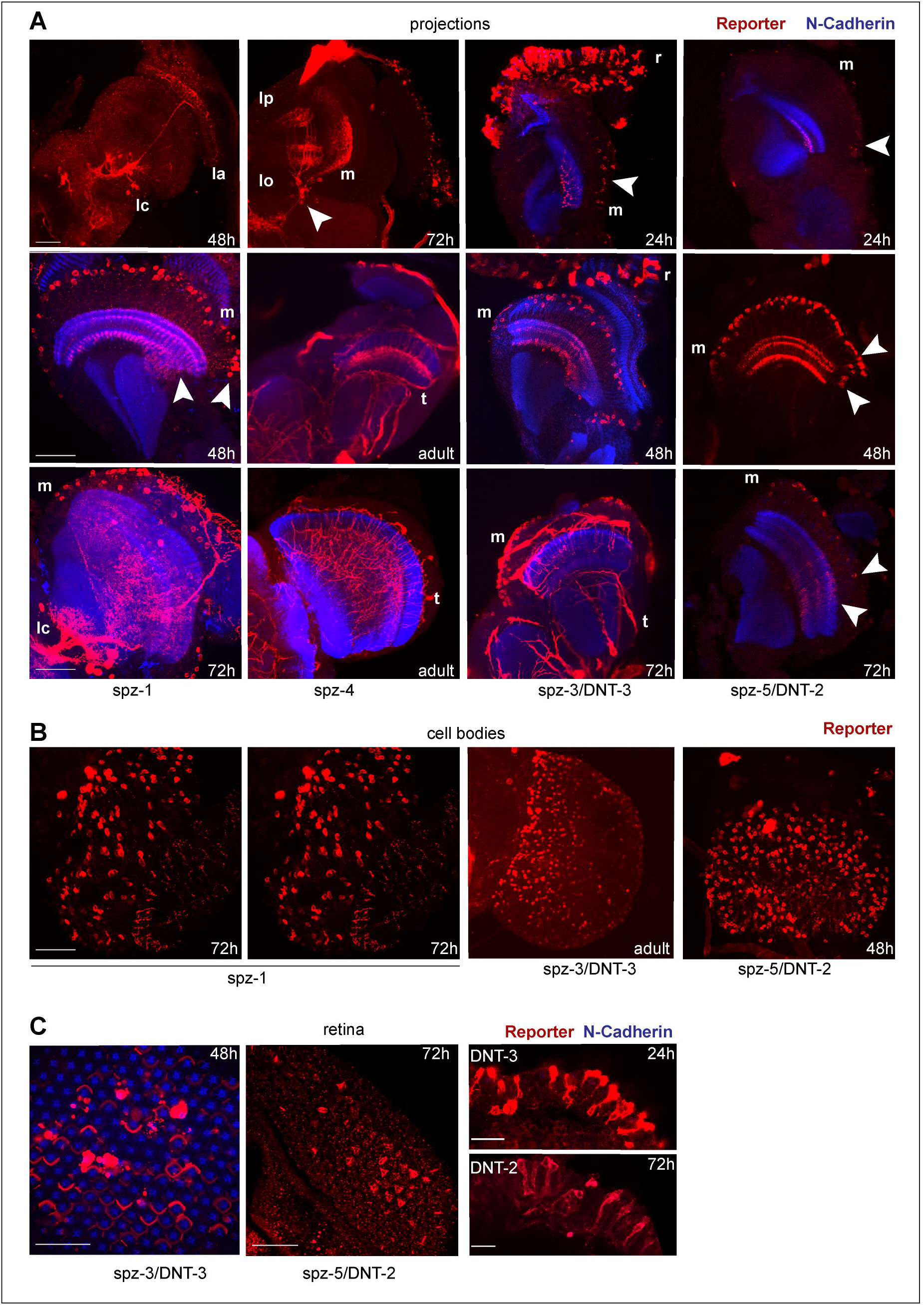
Differential expression of *spz-1, DNT-3 (spz-3), spz-4* and *DNT-2 (spz-5)* during optic lobe development. **(A)** Expression of T2A-Gal4 driving *myrGFP* or *20xUAS6xmCherry* to visualise: *spz-1*^*MIO2318*^*-T2A-Gal4*, in the lobula complex, medulla neurons and projections into lobula and lobula plate; *spz-4*^*MI15678*^*-T2A-GAL4* in the lobula complex and projecting also into medulla, but subsequently prominent in trachea; *spz-3/DNT-3-T2A-Gal4* in retina and medulla, followed by expression in trachea and possibly in glia; *spz-5/DNT-2T2A-Gal4* in medulla neurons, possibly Mi1. **(B)** The location of the cell bodies labelled with *20xUAS6xmCherry* confirms expression for *spz-1* in lobula complex and medulla, and for *spz-3/DNT-3-T2A-Gal4* and *DNT-2-T2A-Gal4* in medulla. **(C)** *spz-3/DNT-3-T2A-Gal4>20xUAS6xmCherry* and *spz-5/DNT-2-T2A-Gal4>MCFO* (anti-HA) are expressed in distinct non-neuronal cells in the retina. Stages are indicated on images, hours after puparium formation (APF). Red: anti-GFP on myrGFP or 6xmCherry; blue: anti-N-Cadherin. Abbreviated genotypes top to bottom: **(A)** spz-1[MIO2318]-T2A-Gal4>myrGP, spz-1[MIO2318]--T2A-Gal4>20UAS6xmCherry; spz-4[MI5678]T2A-Gal4>myrGFP; spz-3-T2A-Gal4>20UAS6xmCherry; DNT-2-T2A-Gal4>20UAS6xmCherry. **(B)** spz-1[MIO2318]--T2A-Gal4>20UAS6xmCherry; spz-3-T2A-Gal4>20UAS6xmCherry; DNT-2-T2A-Gal4>20UAS6xmCherry. **(C)** Left: spz-3-T2A-Gal4>20UAS6xmCherry; DNT-2-T2A-Gal4>MCFO; Right: spz-3-T2A-Gal4>20UAS6xmCherry; DNT-2-T2A-Gal4>myrGFP. For further details, see Table S4. Scale bar: **(A**,**B)** 50μm; **(C)** Left: 50μm and right: 20μm.

Spz-1 and DNT-2 are ligands for Toll-1 and Toll-6 receptors, respectively, and genetic interactions evidence indicates that DNT-3 (Spz-3) is a potential ligand for Toll-8 (Ballard et al., 2014). Tolls are expressed in the optic lobes of adult flies (Li et al., 2020). To visualize the distribution of Tolls in the optic lobes during pupal development, we used the GAL4 lines previously described (Li et al., 2020), driving expression of the reporter *myrGFP* (Figure 2A). *Toll-1-T2A>myGFP* (Singh et al., 2025) revealed initial prominent expression in all optic lobe neuropiles, including the retina, and subsequently was prominent in medulla and lobula complex. *Toll-8*^*MD806*^ *>myrGFP* expression was prominent in the lobula complex, included also some medulla neurons and some lamina neurons. *Toll-6*^*MIO2127*^*>myrGFP* (Li et al., 2020) was initially prominent in the lobula complex but included lamina and medulla neurons, and subsequently expression became more prominent in the lamina. *Toll-2*^*PTV*^*>myrGFP* (Li et al., 2020) was initially found in photoreceptors, lamina, medulla and lobula complex neurons, and by 72 hr APF, *Toll-2*^*PTV*^*>myrGFP* expression was prominent in lamina neurons targeting to the medulla. Using MCFO clones as well as myrGFP, we could identify some of the Toll-8+ neurons as Lawf1, feedback neurons projecting from the medulla to the lamina (Figure 2B), and L2 and L4 lamina neurons (Figure 2B, C); Toll-6+ cells to include L3 and L4 lamina neurons (Figure 2B,D); and Toll-2+ neurons as L1 lamina neurons which target to M1 and M5 medulla layers and L3 lamina neurons that project to M3 (Figure 3B,E) (Hakeda-Suzuki and Suzuki, 2014, Behnia et al., 2014).

**Figure 2.**
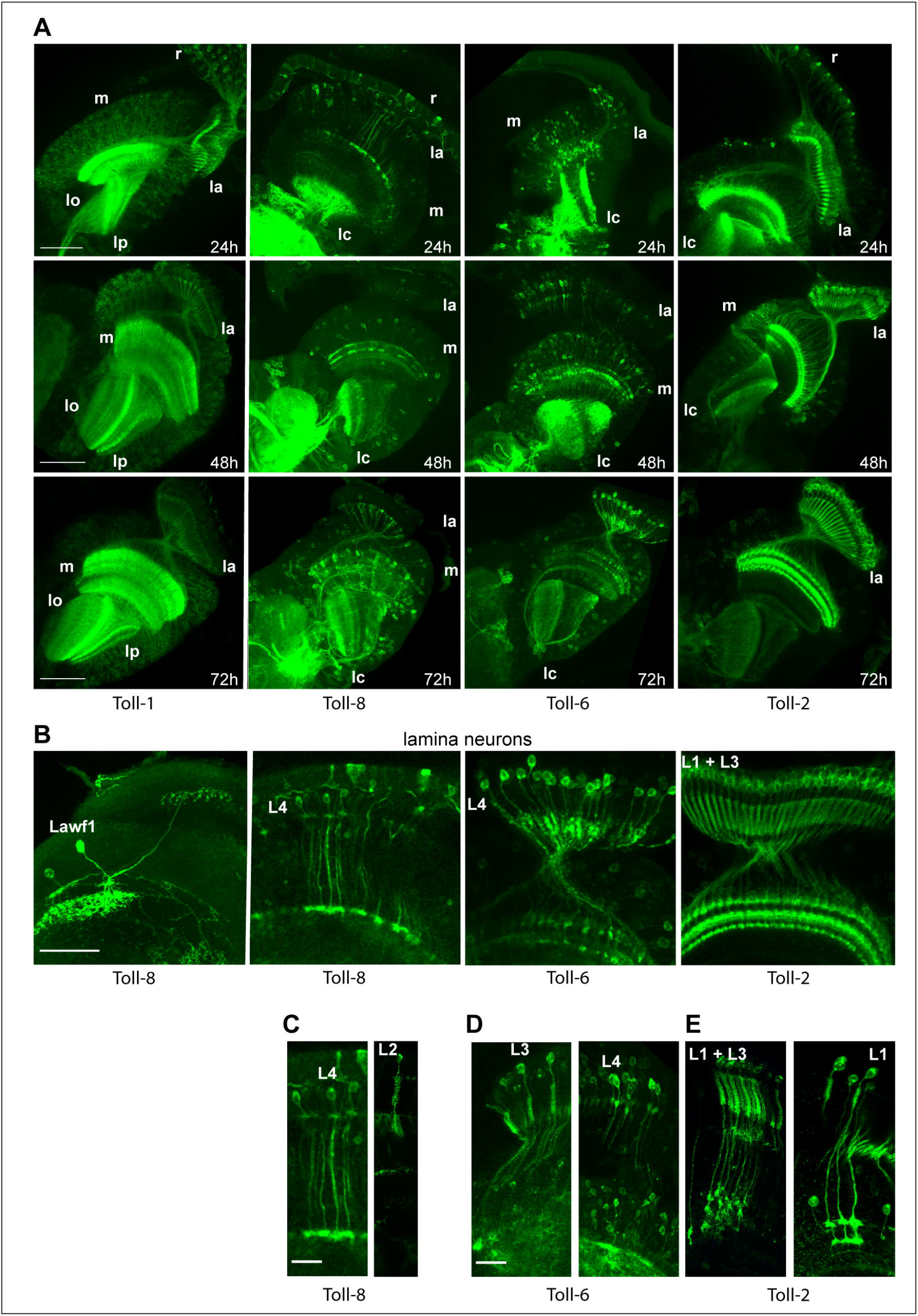
Differential expression of *Toll-1, Toll-2, Toll-6* and *Toll-8* during optic lobe development. **(A)** Expression of *myrGFP* to visualise: *Toll-1-T2A-Gal4* expressing neurons in retina and all optic lobe neuropiles, throughout development; *Toll-8*^*MD806*^*GAL4* predominantly in the lobula complex, plus some lamina and medulla neurons; *Toll-6*^*MIO2127*^*Gal4*, expressed initially more strongly in the lobula complex, but also in lamina and medulla neurons; *Toll-2*^*pTV*^*Gal4* in the retina, most prominently in lamina neurons, but also expressed in some medulla and lobula complex neurons. **(B-E)** MCFO clones with anti-HA antibodies and higher magnification views of myrGFP samples stained with anti-GFP antibodies reveal that Lawf1 medulla and L2 lamina neurons express *Toll-8*, L4 neurons express *Toll-8* and Toll-6, L3 neurons express *Toll-6* and *Toll-2* and L1 neurons express Toll-2. *Toll-2*^*pTV*^*>MCFO*,. Abbreviated genotypes top to bottom: **(A)** *Toll-1-T2A-Gal4>myrGFP; Toll-8*^*MD806*^*GAL4>myrGFP; Toll-6*^*MIO2127*^*>myrGFP; Toll-2*^*pTV*^*>myrGFP*. **(B)** *Toll-8*^*MD806*^*GAL4>MCFO; Toll-8*^*MD806*^*GAL4>myrGFP; Toll-6*^*MIO2127*^*>myrGFP; Toll-2*^*pTV*^*>myrGFP*. **(C)** *Toll-8*^*MD806*^*GAL4>myrGFP;) Toll-8*^*MD806*^*GAL4>MCFO; Toll-6*^*MIO2127*^*>MCFO; Toll-6*^*MIO2127*^*>myrGFP; Toll-2*^*pTV*^*>MCFO*. For further details, see Table S4. Scale bar: **(A**,**B)** 50μm; **(C**,**D)** 20μm.

**Figure 3.**
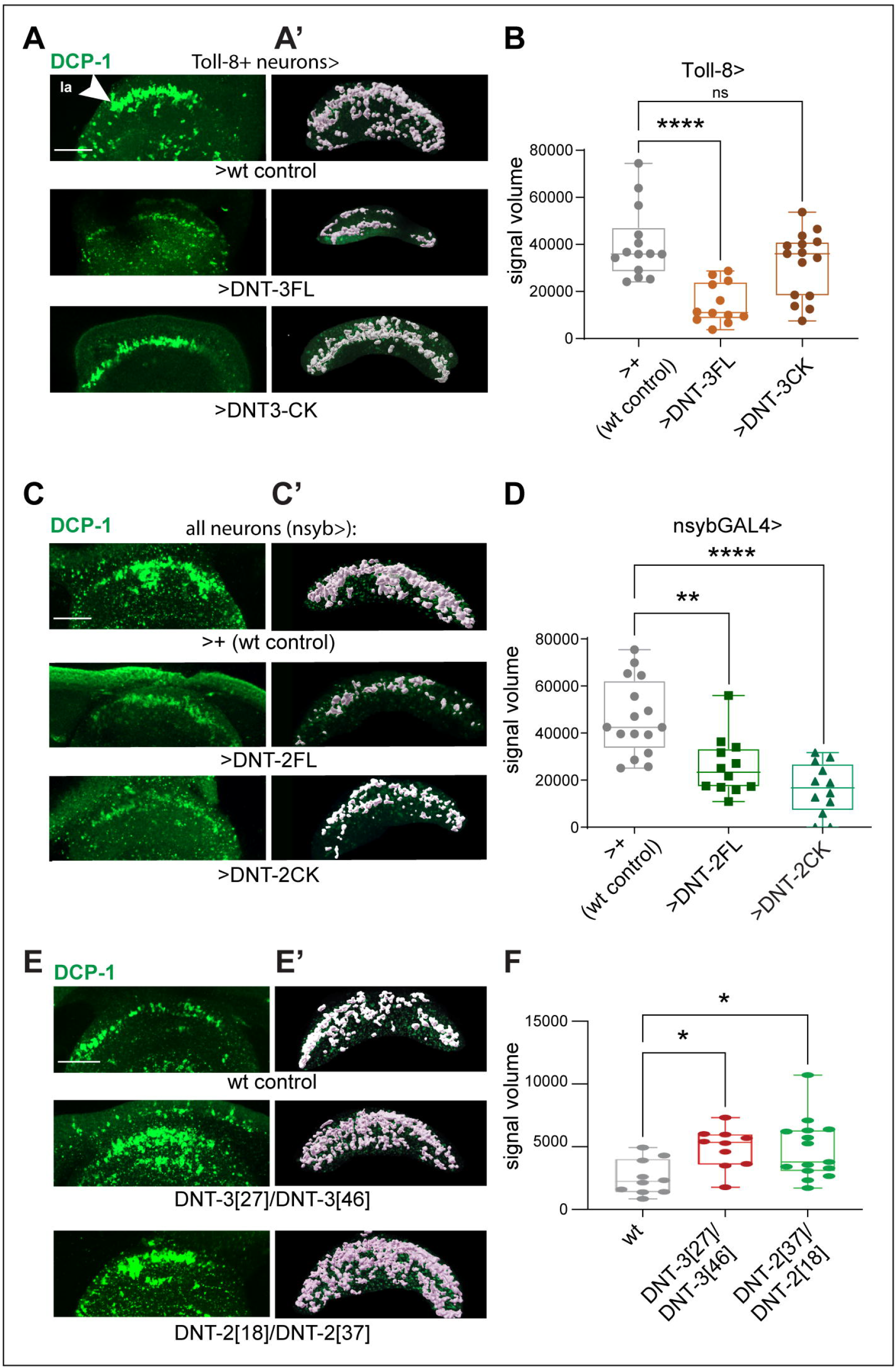
DNT-2 and DNT-3/spz-3/DNT-3 can and are required to maintain cell survival during optic lobe development. **(A-B)** Over-expression of *spz-3/DNT-3* in Toll-8 neurons *(Toll-8>spz-3/DNT-3)* and **(C-D)** of *DNT-2/spz-5* in all neurons *(nsyb>DNT-2)* can rescue naturally occurring cell death, in the lamina. Apoptotic cells labelled with anti-Dcp1 at 24h APF. **(A-B)** ONE WAY ANOVA ****p<0.0001, and Dunnett’s multiple comparisons correction to a fixed control. **(C-D)** ONE WAY ANOVA ****p<0.0001, and Dunnett’s multiple comparisons correction to a fixed control. **(E-F)** *spz-3*^*46*^*/spz-3*^*27*^ (i.e. hereby renamed DNT-3 mutants) and *DNT-2*^*18*^*/DNT-2*^*37*^ (i.e. also called *spz-5*^*18*^*/spz-5*^*37*^) trans-heterozygous loss of function mutants had increased apoptosis at 24h APF. ONE WAY ANOVA *p<0.05, and Dunnett’s multiple comparisons correction to a fixed control. Images in **(A’, C’, E’)** are Imaris 3D-renderings of lamina ROI used for signal volume measurements. FL: full length; -CK: mature, cleaved, cystine-knot domain. Asterisks indicate multiple comparisons corrections: *p<0.05, **p<0.01, ***p<0.001, ****p<0.0001. For further details, see Table S4. Scale bar: 50μm.

We compared our reporter-based profiles with published scRNAseq datasets of the optic lobe through development (Kurmangaliyev et al., 2020, Ozel et al., 2021) (Supplementary Figures S1-7). Consistently with cell biology data, in scRNAseq spz ligand expression was low and dynamic, and *Toll* receptors were expressed more abundantly and more widely. Overall, the expression in the scRNAseq dataset (Kurmangaliyev et al., 2020) of the *spz* ligands and *Toll-8* (also known as Tollo) data were less consistent with the cell biology data, whereas the expression of *Toll-1, -2* and *-6* confirmed cells seen with the cell-biology based reporters. Most particularly, *Toll-2* mRNA (synonym 18w) was found in L1 and L3 lamina neurons over time, plus also in L5 at 24h APF, and *Toll-6* mRNA was found in L2, L3, L4 over time, plus also in L1 at 24h (Supplementary Figure 1-6). Focusing on *DNT-2 (spz-5)* and *Toll-2 (18w), DNT-2* mRNA was only found in Mi1 neurons in low levels at 48h and instead was found in high levels in clusters (some unclassified) which could include other medulla neurons (Supplementary Figure S7). Between 24-48h APF, *Toll-2* was highly expressed in L1 and L3 lamina neurons, consistently with reporter-based data (Figure 2), as well as in L4 and L5, and in medulla neurons including Dm3, Dm9, Mi1 and in low levels in Tm3 (Supplementary Figure S7). From 72h APF, onwards, *Toll-2* expression decreased in L1, Tm3 and Dm9, and it increased in L3 and T4 and T5 neurons. Importantly, this showed that *Toll-2* was expressed in connecting cells (L1, Mi1, Tm3, Dm9, T4) that could receive DNT-2 ligand secreted from medulla neurons (Supplementary Figure S7).

To conclude, at the time of naturally occurring cell death (0-48h APF), *Toll-1* is highly expressed throughout the optic lobe; *Toll-2, -6* and *-8* are expressed in the medulla; *Toll-8* and *Toll-6* are prominently expressed in the lobula complex, and *Toll-6* and *Toll-2* are prominent in the lamina. Interestingly, each lamina neuron type expresses distinct Tolls, or combination of Tolls: L1 neurons express *Toll-2* (Figure 2B,E); L2 express *Toll-6* and possibly *Toll-8* (Figure 2B,C and scRNAseq); L3 express *Toll-2* and *Toll-6* (Figure 2D,E); and L4 express *Toll-6* and possibly *Toll-8* (Figure 2C,D). Remarkably, L1 neurons receive input from R1-R6, project to the medulla making synaptic contact with Mi1 medulla neurons at the M1 layer and *Toll-2* is expressed in L1 neurons and *DNT-2* in Mi1 (or related) medulla neurons. This suggests that the secreted ligand DNT-2 could reach the Toll-2 receptor.

The neurotrophins DNT-2 and DNT-3 promote neuronal survival during optic lobe development

DNT-2 (Spz-5) promotes cell survival in the embryonic central nervous system (CNS) and in adult brains, DNT-1 (Spz-2) promotes cell survival in the embryonic CNS, and DNT-3 (Spz-3) promotes cell survival in the adult brain; and both DNT-2 and DNT-3 also have neurotrophin functions at the neuro-muscular junction (Zhu et al., 2008, Coutinho-Budd et al., 2017, Sun et al., 2024, Ballard et al., 2014, Ulian-Benitez et al., 2017). To ask whether *DNTs* can promote cell survival in developing optic lobes, we over-expressed *DNT-2 (spz-5)* and *DNT-3 (spz-3*) and visualized dying cells with the apoptotic marker anti-Dcp1 at the peak of naturally occurring cell death (24h APF). Spz family ligands, like mammalian neurotrophins, can be found in full-length or cleaved forms consisting of the evolutionarily conserved neurotrophin cystine-knot domain (Foldi et al., 2017, Zhu et al., 2008, DeLotto and DeLotto, 1998, Hu et al., 2004). Over-expressed cystine-knot forms from transgenic flies may not be secreted, but can be functional upon in vivo over-expression, by acting cell-autonomously (Foldi et al., 2017, Sun et al., 2024, Zhu et al., 2008, Ulian-Benitez et al., 2017, Hu et al., 2004). By contrast, over-expressed full-length forms of *DNT-2* and *DNT-3* are spontaneously cleaved and secreted both in cell culture and in vivo, thus enabling non-autonomous activation of their receptors (McIlroy et al., 2013, Foldi et al., 2017, Sun et al., 2024, Coutinho-Budd et al., 2017). Both *DNT-2* and *DNT-3* are expressed in medulla neurons (Figure 1), which could influence non-autonomously neurons in the lamina, medulla and lobula complex. Thus, we focused on *DNT-2* and *DNT-3* and tested their functions in the optic lobe.

As Toll-8 is the proposed receptor for DNT-3 (Spz-3) (Ballard et al., 2014), we over-expressed *DNT-3* in Toll-8+ neurons. Over-expression of *DNT-3-full length (DNT-3FL)* but not cleaved *DNT-3* cystine-knot *(DNT-3 CK)* in Toll-8+ cells (i.e. *Toll-8>DNT-3FL or Toll-8>DNT-3CK*, respectively) reduced the incidence of Dcp1+ apoptosis in the lamina (Figure 3A-B). *DNT-3FL* is secreted (Coutinho-Budd et al., 2017) and thus can reach broadly, whereas presumably *DNT-3CK* is not (see Discussion), and it might be unable to rescue cell survival cell-autonomously due to the weak and irregular expression of *Toll-8* in the lamina with this Gal4 line (see Figure 2). Over-expression of *DNT-3FL* also reduced Dcp1+ incidence outside the lamina, throughout the medulla and lobula complex (Supplementary Figure S9A,B). To test whether DNT-2 could promote cell survival during optic lobe development, we over-expressed full-length *DNT-2FL* or cleaved *DNT-2CK* in all neurons with *nsybGAL4*. This reduced the incidence of Dcp1+ apoptosis in the lamina and outside the lamina too (Figure 3C-D and Supplementary Figure S9C,D). Altogether, secreted DNT-2 and -3 could promote cell survival during optic lobe development, preventing naturally occurring cell death. Conversely, we asked whether DNT-2 and -3 are required to maintain cell survival during optic lobe development. We generated *DNT-3 (spz-3)* loss of function mutants by P-element mobilization. *DNT-2* and *DNT-3* loss of function mutants caused considerable cell debris in the medulla and lobula complex, which compromised the analysis in this region, so we focused on the lamina. We found that in *DNT-3*^*46*^*/DNT-3*^*27*^ (i.e. *spz-3*^*46*^*/spz-3*^*27*^) pupae, Dcp1 increased in the lamina compared to wild-type controls (Figure 3E-F). Similarly, in null *DNT-2*^*18*^*/DNT-2*^*37*^ mutants (Foldi et al., 2017, Sun et al., 2024, Ulian-Benitez et al., 2017) Dcp-1 also increased in the lamina (Figure 3E-F). These data show that both DNT-2 and DNT-3 are required to maintain cell survival of lamina cells during optic lobe development.

To conclude, *DNT-3 (spz-3)* and *DNT-2 (szp-5)* can rescue naturally occurring cell death and are required to maintain cell survival in the lamina during optic lobe development. Furthermore, the data suggest their effect could be broader as they can also promote cell survival outside the lamina, in the optic lobe.

### DNT-2 promotes neuronal survival via Toll-2 in optic lobe development

To test whether the ability of DNTs to promote cell survival in the optic lobe depends on signalling through their Toll receptors, we focused on DNT-2. DNT-2 binds promiscuously Toll-6 and Toll-7 (McIlroy et al., 2013). Both Toll-6 and Toll-2 are expressed in lamina neurons (Figure 2). Toll-2 is neuroprotective (Li et al., 2020), but its activating ligand was unknown. Remarkably, *DNT-2* is expressed in Mi1 medulla neurons (Figure 1A), which connect to L1 neurons (Nern et al., 2025) which express *Toll-2* (Figure 2A,B,E) (and Supplementary Figures 1-7, scRNAseq data from (Kurmangaliyev et al., 2020)). *Toll-2*^*pT*V^GAL4 flies are heterozygous mutant for *Toll-2*, and, remarkably, in combination with *DNT-2* homozygous mutants resulted in semi-lethality, revealing a functional interaction between these two genes. This strongly suggested that Mi1 medulla neurons could secrete DNT-2, which could bind Toll-2 in L1 lamina neurons. We tested this using genetic epistasis analysis. First, we found that RNAi knock-down of *Toll-2* in all neurons (i.e. *nybGAL4>Toll-2RNAi*^*GD36305*^) increased Dcp1+ apoptosis in the lamina (Figure 4A,B), meaning that Toll-2 is required to maintain lamina neuron survival, consistently with mediating the functions of a DNT ligand. Importantly, *Toll-2RNAi*^*GD36305*^ knock-down prevented over-expressed DNT-2FL from rescuing cells from apoptosis, as Dcp1+ levels remained high in this genotype (Figure 4A,B). These data demonstrate that DNT-2 functions via Toll-2 to promote lamina neuron survival.

**Figure 4.**
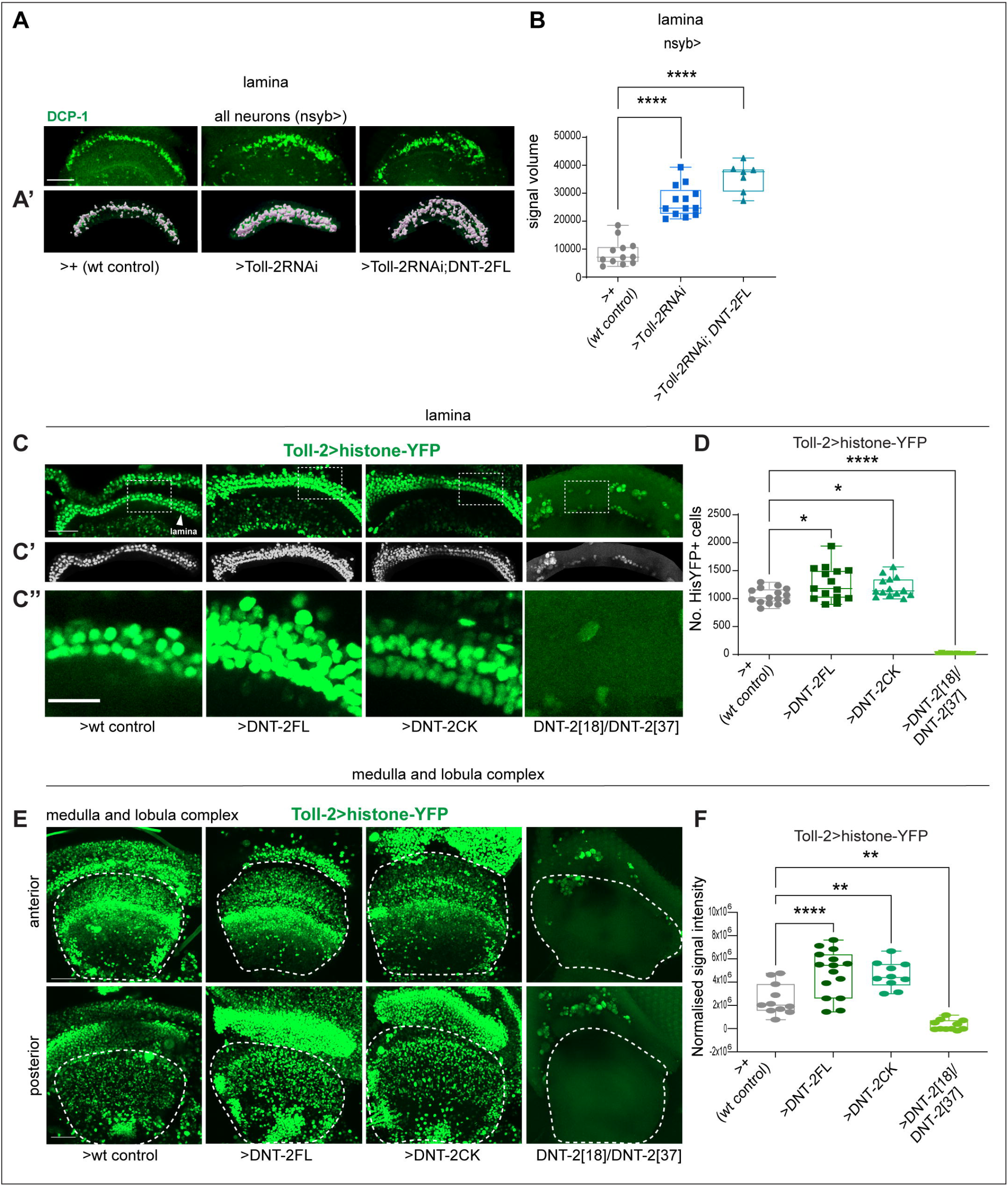
DNT-2 functions with Toll-2 to regulate cell survival in the optic lobe. **(A**,**B)** Pan-neuronal RNAi knock-down of *Toll-2 (i.e. nsybGAL4>UAS-Toll-2*^*GD36305*^*-RNAi)* induced apoptosis, and the rescue of naturally occurring cell death by DNT-2 gain of function was prevented when *Toll-2* was also knocked-down (i.e. epistasis: *nsybGAL4>UAS-Toll-2* ^*GD36305*^*-RNAi, DNT-2FL*). 3D-measurement of signal volume with Imaris. Kruskal-Wallis ANOVA ****p<0.0001, and Dunn’s multiple comparisons correction to a fixed control. **(C**,**D)** In the lamina, over-expression of *DNT-2FL* or *DNT-2CK* in Toll-2 cells increased cell number *(Toll-2>his-YFP, DNT-2FL* and *Toll-2>his-YFP, DNT-2CK)*, but Toll-2> HisYFP+ cells were virtually all lost in *DNT-2*^*18*^*/DNT-2*^*37*^ mutants. Automatic cell counting with DeadEasy Optic Lobe. Welch ANOVA ****p<0.0001, and Dunn’s multiple comparisons correction to a fixed control. **(E**,**F)** Outside the lamina, in the medulla and lobula complex, over-expression of *DNT-2FL* or *DNT-2CK* in *Toll-2>hisYFP* cells increased signal intensity, and virtually all *Toll-2>his-YTP+* signal was lost in *DNT-2*^*18*^*/DNT-2*^*37*^ mutants, with only HisYFP+ macrophages remaining. ONE WAY ANOVA ****p<0.0001, and Dunnett’s multiple comparisons correction to a fixed control. Projections in **(E**,**F)** of anterior optic lobes correspond approximately to the medulla, and posterior correspond approximately to the lobula complex; signal intensity was measured using Fiji and normalized to background signal. Dotted lines in **(C)** indicate region shown in **(C’’)** magnified; in **(E)** indicate ROI excluding the lamina used for signal intensity measurements. FL: full length; -CK: cystine-knot domain. Asterisks indicate multiple comparisons correction tests: *p<0.05, **p<0.01, ***p<0.001, ****p<0.0001. For further details, see Table S4. Scale bar: **(A**,**C**,**E)** 50μm; **(C’’)** 20μm.

Apoptosis can be verified by testing whether changes in cell death lead to changes in cell number. Toll-2+ lamina neurons are clearly distinct when labelled with the nuclear reporter histone-YFP (i.e. *Toll-2>hisYFP*) and could be quantified automatically using DeadEasy. Thus, we asked whether altering DNT-2 levels could result in changes in Toll-2>hisYFP+ lamina cell number. We found that the reduction in apoptosis caused by DNT-2FL and DNT-2CK gain of function led to an increase in Toll-2>hisYFP+ lamina neuron number compared to wild type controls (Figure 4C,D). Conversely, the increase in apoptosis we had seen in *DNT-2*^*18*^*/DNT-2*^*37*^ null mutants resulted in the dramatic loss of virtually all Toll-2>hisYFP+ lamina cells (Figure 4C,D). Macrophages loaded with HisYFP and distributed mostly between the retina and lamina could be observed across these samples (Figure 4C), suggesting they had engulfed dead cells. Finally, we asked whether altering DNT-2 levels might also affect the number of Toll-2+ cells outside the lamina, throughout the medulla and lobula complex. As too many cells were labelled with Toll-2>HisYFP, automatic cell counting was not possible and we measured signal intensity instead. This showed that over-expression of either *DNT-2FL* or *DNT-2CK* increased Toll-2>HisYFP cell signal intensity in the medulla and lobula complex (Figure 4E,F), consistently with an increase in cell number. Most dramatically, *DNT-2*^*18*^*/DNT-2*^*37*^ null mutants revealed a complete loss of Toll-2+ cells also outside the lamina, in the medulla and lobula complex (Figure 4E,F). These data demonstrate that the decrease in apoptosis caused by DNT-2 gain of function increased the number of Toll-2+ cells, and the increase in apoptosis in *DNT-2* mutants caused Toll-2+ cell loss.

Together, these data show that DNT-2 functions as a ligand for Toll-2 to maintain the survival of neurons in the lamina, medulla and lobula complex during optic lobe development.

### DNT-2 and Toll-2 regulate targeting of lamina neurons to the medulla

In the vertebrate nervous system, neurotrophic factors are produced in limiting amounts, and only neurons that receive them when in proximity to target neurons are spared from naturally occurring cell death (Davies, 2003, Levi-Montalcini, 1987, Lu et al., 2005). In this way, cell number is adjusted during connectivity, enabling the establishment of appropriate, functional neural circuits (Davies, 2003, Levi-Montalcini, 1987, Lu et al., 2005). In the *Drosophila* pupa, connectivity of lamina to medulla neurons takes place at 30-48h APF, and between medulla and lobula complex at 60-70h APF (Kurmangaliyev et al., 2020, Millard and Pecot, 2018, Pecot et al., 2014, Hadjieconomou et al., 2011). Importantly, the expression of synaptic markers starts at 24h, peaks at 60h APF and spontaneous neuronal activity takes place at 48h APF, meaning that at least some neural circuits are already connected by this point (Kurmangaliyev et al., 2020). Thus, the period of naturally occurring cell death overlaps with connectivity. To ask whether DNT-2 with Toll-2 could function during neural circuit development, we tested whether DNT-2 is required by lamina neurons for connecting to target neurons in the medulla. As *Toll-2* is expressed in L1 and L3 lamina neurons (Figure 2E, MCFO), to visualize L1 neurons we used L1-specific split-GAL4 (R48A08AD; R66A01DBD) (Tuthill et al., 2013) to drive the expression of *mCD8-GFP* or *myr-GFP* reporters, and tested the effect of altering gene expression on axonal and dendritic patterns. L1 neurons normally project along columns that can be labelled with mAb24B10, and target to layers M1 and M5 of the medulla. The M1 connection is particularly relevant as here L1 neurons (which express Toll-2) connect with Mi1 medulla neurons (which express DNT-2). When we overexpressed *DNT-2* in L1 neurons (i.e. *L1-splitGAL4>mCD8GFP, DNT-2FL or L1-splitGAL4> mCD8GFP, DNT-2CK*), L1 axonal terminals in the M1 layer frequently misrouted, crossing over to neighboring 24B10+ columns (Figure 5A,C). To test whether Toll-2 was required for appropriate connectivity, we knocked down *Toll-2* using RNAi (i.e. *L1-split-GAL4>myrGFP, Toll-2RNAi*^*GD36305*^) and this produced two phenotypes: a milder (50% penetrance n=10 10 brains) and a severe (50% penetrance n=10 brains) phenotype. In severely affected optic lobes, most L1 neurons were missing, consistently with the increase in cell death (Figure 5), and surviving neurons could misroute at M1 layer to nearby columns (Figures 5B,D). In mildly affected optic lobes, more neurons remained, and L1 axonal terminals frequently misrouted (Figures 5B,D). Interestingly, *Toll-2RNAi* knock-down did not alter the phenotype caused by *DNT-2FL* overexpression, and impaired targeting to the same extent as each genetic manipulation alone (Figures 5B,D). These data showed that both excess and deficit in DNT-2, and loss of Toll-2 impaired axonal targeting. This is consistent with the notion that neurotrophic factors are released in limited amounts and both up- or down-regulation in their levels disrupt the information carried by secreted ligand gradients (Zhu et al., 2008, Davies, 2003).

**Figure 5.**
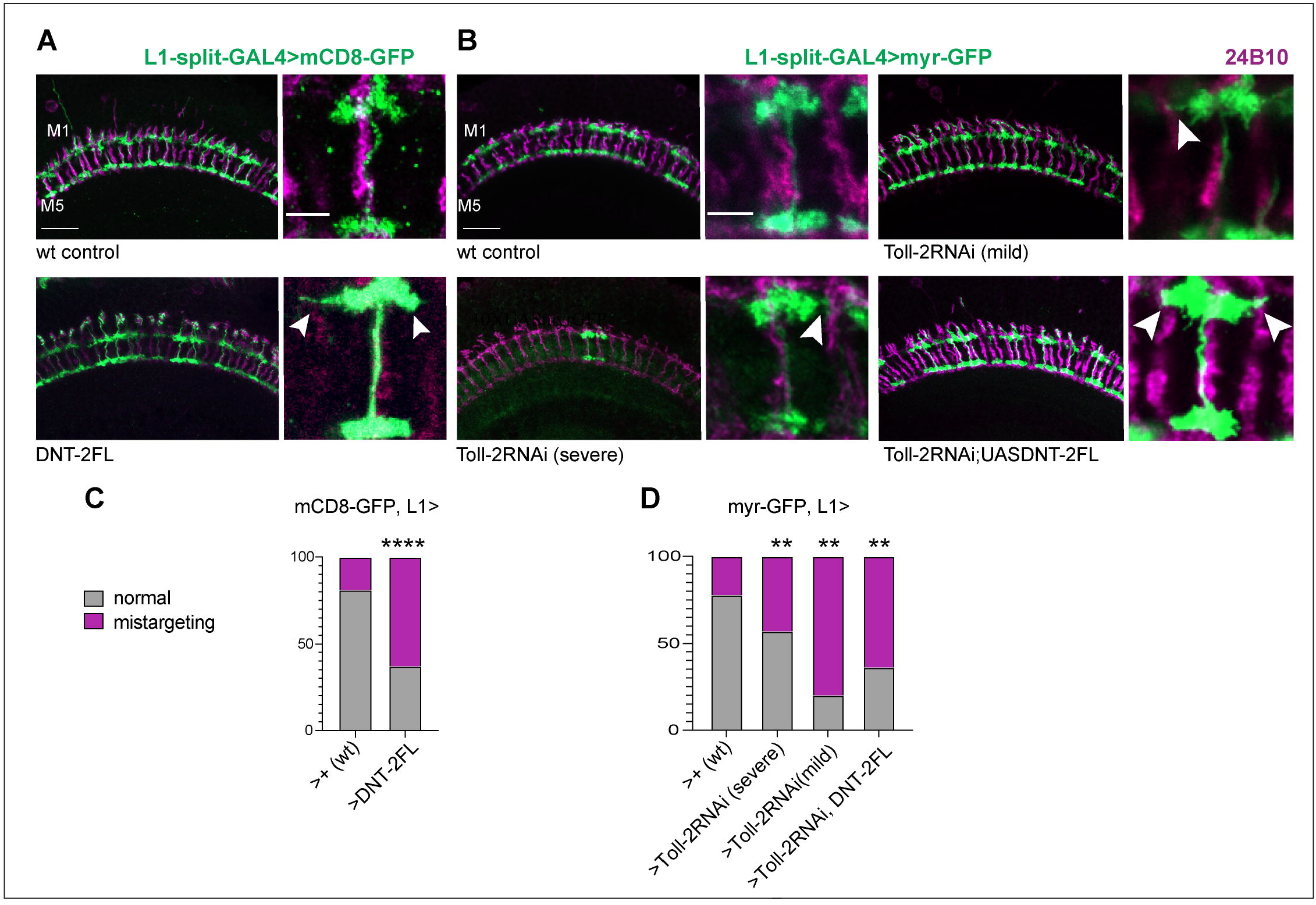
DNT-2 with Toll-2 regulate connectivity at the medulla. **(A)** Over-expression of DNT-2FL in L1 lamina neurons *(L1 split>DNT-2FL)* caused axonal misrouting at M1 layer (arrowheads); quantification in **(C). (B)** *L1 split>Toll-2*^*GD36305*^*-RNAi* knockdown in L1 lamina neurons caused a severe phenotype in which most L1 neurons were missing (50% penetrance, n=10 number of brains), a mild phenotype (50% penetrance, n=10), and remaining neurons misrouted; quantification in **(D)**. *Toll-2*^*GD36305*^*-RNAi* knock-down together with DNT-2 over-expression *(L1 split>Toll-2*^*GD36305*^*-RNAi, DNT-2FL)* also caused misrouting. FL: full length. Fisher’s Exact test **p<0.01, and post-hoc Bonferroni multiple comparisons corrections. Asterisks refer to multiple comparisons corrections **p<0.01, ****p<0.0001. For further details, see Table S4. Scale bar: **(A**,**B)** Left: 20μm. Right: 5μm.

To conclude, these data show that DNT-2 and Toll-2 are required for appropriate connectivity of L1 neurons to target Mi1 medulla neurons at M1 medulla layer.

### DNT-2 regulates lamina neuron dendritic complexity via Toll-2

Lamina neurons have a distinctive dendritic morphology, can have mushroom shaped dendritic spines, and are known to be plastic (Weber et al., 2009). To investigate whether DNT-2 and Toll-2 may affect L1 neuron dendritic morphology, we used the *L1-split-GAL4* driver line (R48A08AD; R66A01DBD) (Tuthill et al., 2013), as above, and visualized L1 dendrites with *UASmCD8-GFP* or UAS-myrGFP and traced the dendritic shape and spines in 3D. We overexpressed *DNT-2FL (i.e. L1-splitGAL4>mCD8GFP, DNT-2FL)* in L1 neurons, and this resulted in a significant increase in dendritic volume compared to wild-type controls, and larger and more abundant dendritic spines (Figure 6A,C). To investigate whether Toll-2 was involved, we knocked down *Toll-2* expression with RNAi (i.e. *L1-split-GAL4>myrGFP, Toll-2RNAi*^*GD36305*^). As before, this resulted in some optic lobes with very few L1 neurons, as most of them were missing (“severe”, penetrance 60% n=5 brains) and other optic lobes that had multiple L1 neurons (“mild”, penetrance 40% n=5 brains). *Toll-2* RNAi knockdown resulted in a decrease in overall dendrite size, and a marked decrease in dendritic spines (Figure 6B,D). This suggests that Toll-2 is essential for maintaining both the size and branching architecture of L1 dendrites. Finally, *Toll-2 RNAi* knock-down prevented the increase in dendritic size that would have been caused by *DNT-2FL* over-expression (i.e., *L1-split-GAL4>myrGFP, Toll-2RNAi*^*GD36305*^, *DNT-2FL*), demonstrating that DNT-2 functions via Toll-2 to regulate lamina dendrite size and complexity (Figure 6B,D).

**Figure 6.**
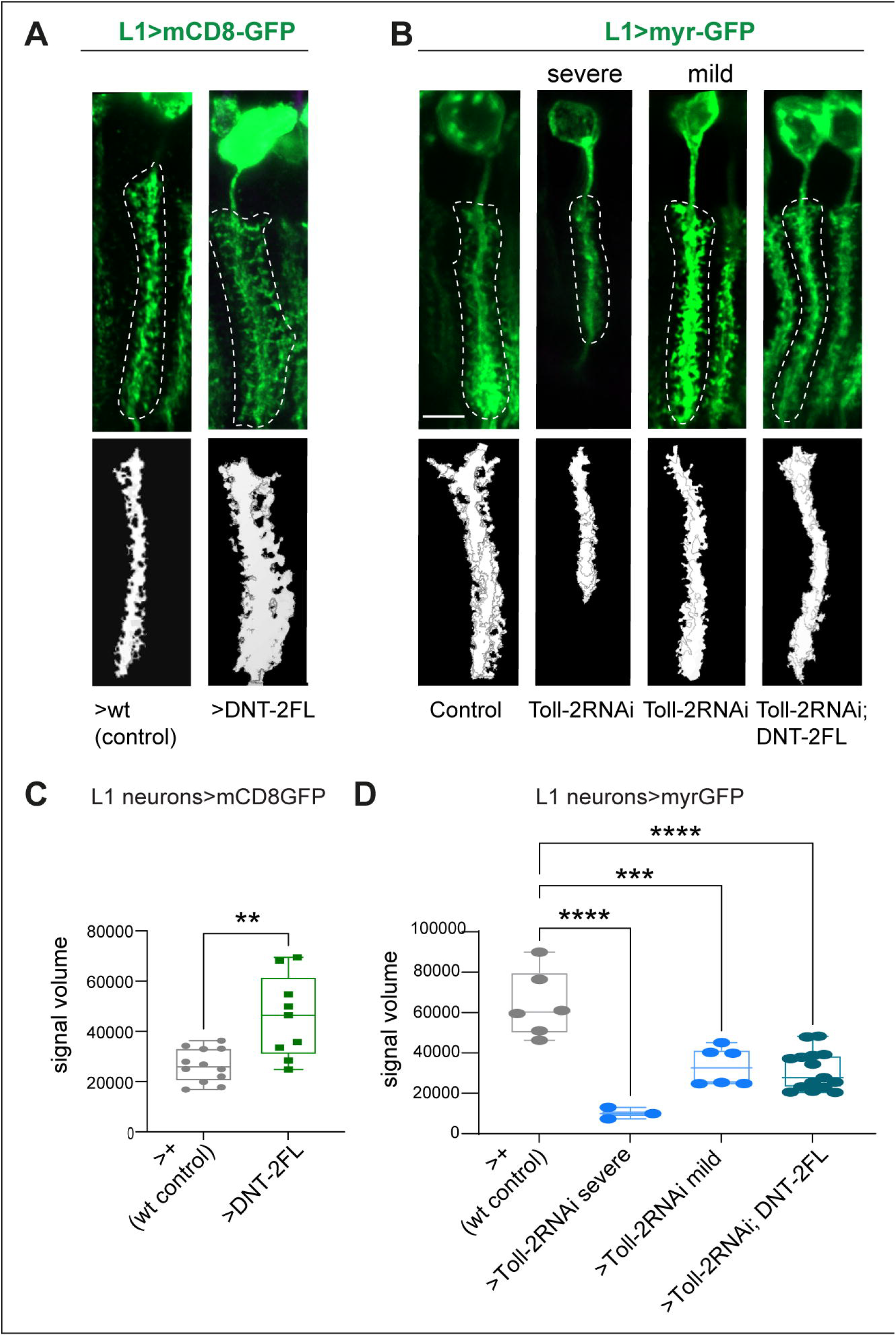
DNT-2 with Toll-2 can modify lamina dendrite size. **(A)** Over-expression of DNT-2 in L1 neurons increased dendrite size *(L1 split>mCD8-GFP, DNT-2FL)*. **(C)** Quantification using Amira. Student t test p<0.01. **(B)** *Toll-2*^*GD36305*^*-RNAi* knockdown decreased dendrite size: in severely (40% penetrance n=5) and mildly (60% penetrance n=5) affected specimens *(L1 split>myrGFP, Toll-2*^*GD36305*^*-RNAi). Toll-2RNAi* ^*GD3630*^ knock-down rescued the increase in dendrite size by *DNT-2FL* over-expression *(L1 split>myrGFP, Toll-2*^*GD36305*^*-RNAi, DNT-2FL)*. **(D)** Quantification of data shown in **(B)**: Kruskal Wallis ANOVA **p<0.01, and Dunn’s multiple comparisons correction to a fixed control. **(A, A’**,**B, B’)** Top images show raw images with a dotted line around the dendrites; bottom images show Amira 3D rendering of dendrite volume. FL: full length. Asterisks in **(D)** refer to multiple comparisons corrections: **p<0.01 ***p<0.001, ****p<0.0001. For further details, see Table S4. Scale bars: 5μm.

## Discussion

Neural circuits develop from a highly reproducible choreography of cellular events that reveals the robustness of development. Visual system development in *Drosophila* is stereotypic and over time this enables 750-800 neuron-hosting units to form precise connections across lamina cartridges, medulla columns and layers, preserving retinotopy and enabling appropriate vision. The reproducibility and robustness of circuit development emerges from the combination of multiple underlying mechanisms. Genetic cascades control the extent of neural stem cell proliferation, short and long-range molecules aid axonal navigation and glial migration, cell-cell interactions drive reassortment and superposition of axons, and molecular tags between input and target enable synaptic matching (Agi et al., 2024, Courgeon and Desplan, 2019a, Holguera and Desplan, 2018, Malin et al., 2024, Pecot et al., 2014, Melnattur and Lee, 2011, Millard and Pecot, 2018). Concomitantly, a wave of cell death takes place in *Drosophila* optic lobe development (Hara et al., 2013, Hara et al., 2018, Togane et al., 2012). Such cell death could reflect the process of metamorphosis, loss of abnormal cells and adjustments between interacting cell populations, including during the emergence of neural circuits (Hara et al., 2013, Pinto-Teixeira et al., 2016, Togane et al., 2012, Pecot et al., 2014). In fact, throughout animal development, between 50% (e.g. in *Drosophila*) and 80% (e.g. in vertebrates) are lost to naturally occurring cell death (Rogulja-Ortmann et al., 2007, Pinto-Teixeira et al., 2016, Davies, 2003, Dekkers et al., 2013, Raff et al., 1993). The control of cell survival during periods of naturally occurring cell death enables adjustments of neurons to their targets and connecting neurons, between neuronal and glial cell populations, and facilitates the establishment of neural circuitry conducive to appropriate behaviour. Our findings demonstrate that *DNT-2 (i.e. spz-5)* and *DNT-3 (i.e. spz-3)* have evolutionarily conserved functions maintaining neuronal survival during naturally occurring cell death in visual system development of *Drosophila*. Furthermore, *DNT-2* produced from medulla neurons maintains the survival, targeting and dendritic complexity of connecting L1 neurons that express the Toll-2 receptor. These data support the notion that neurotrophism is a fundamental property of nervous system development to enable adjustments between interacting cell populations.

We have shown that *DNT-2 (spz-5)* and *DNT-3 (spz-3*) are expressed in the retina and medulla and are required for and can promote survival of lamina and medulla neurons. Analysis of published scRNAseq data (Kurmangaliyev et al., 2020) confirmed the expression profiles of *Toll-1, -2* and *-6*, as observed with the GAL4/UAS reporters, whereas evidence for *Toll-8* was variable in the microscopy data, and ligands revealed clearer profiles with reporters. The discrepancies between the reporter-based and the scRNAseq profiles could reflect different sensitivity (e.g. GAL4/UAS creates an amplification) and temporal resolution (i.e. microscopy data reveal the perdurance of the reporters, whereas scRNA is a snapshot in time in gene expression). Importantly, our data show consistent relationships between the microscopy expression profiles and loss of function phenotypes.

Consistently with these findings, we have shown that the survival of L1 neurons depends on DNT-2 functioning together with Toll-2. DNT-2 is a well-known ligand for Toll-6 and Kek-6, but it can also bind promiscuously Toll-7 and Kek-2 (McIlroy et al., 2013, Ulian-Benitez et al., 2017). Our data here strongly indicate that DNT-2 is also a ligand for Toll-2: *Toll-2* is expressed in L1 neurons that connect to Mi1 neurons expressing *DNT-2*; loss of function for *DNT-2* or *Toll-2* resulted in an increase in Dcp1 levels in the lamina; trans-heterozygosis for *DNT-2* and *Toll-2* caused semi-lethality; the rescue of cell death caused by *DNT-2* over-expression could be prevented by knocking-down *Toll-2*; the increase in L1 dendrite size caused by *DNT-2* over-expression could be rescued by knocking-down *Toll-2*. These data demonstrate that *DNT-2* and Toll-2 function together in visual system development.

We have shown that over-expression of *DNT-3* and *DNT-2* can both rescue naturally occurring cell death in the lamina and also outside the lamina (i.e. in the medulla plus lobula complex), that *DNT-2, Toll-2* and *DNT-3* are required for lamina neuron survival, and that loss of function for *DNT-2* in a heterozygous *Toll-2* mutant background results in the virtually complete loss of Toll-2+ neurons in the lamina, medulla and lobula complex. Our findings are consistent with a previous report that *DNT-3 (spz-3)* functions as a neurotrophin and promotes cell survival in the adult central brain (Coutinho-Budd et al., 2017). *DNT-2* has also previously been shown to promote cell survival in the embryonic CNS and adult brain, and its receptor *Toll-6* in the larval and pupal ventral nerve cord (Foldi et al., 2017, McIlroy et al., 2013, Sun et al., 2024, Zhu et al., 2008). In summary, our data show that during the peak of apoptosis sweeping across the optic lobe in pupal development, DNT-2 and DNT-3 are required for and can promote lamina neuron survival, and strongly suggest that they also regulate survival of medulla and lobula complex neurons. Our data suggest that at least DNT-3 (Spz-3) could also regulate the survival of other lamina neurons with another Toll receptor. They also suggest that survival of different lamina neurons might depend on different Tolls or combinations of Tolls. In fact, L2, L3 and L4 neurons express *Toll-2, Toll-6* or *Toll-*8 receptors that could also maintain their survival, perhaps with other ligands or combination of ligands. In fact, loss of function for *DNT-3 (spz-3)* causes lamina cell death that is not naturally compensated for by DNT-2. Finally, interference with the normal levels of DNT-2 and Toll-2 also impaired axon targeting and dendritic morphology, consistently with the coupling between cell survival with connectivity.

DNTs can function both as full-length and cleaved, mature ligands. Full length and cleaved forms for DNT-1 (Spz-2), DNT-2 (Spz-5) and DNT-3 (Spz-3) can be found naturally in the brain (Coutinho-Budd et al., 2017, Foldi et al., 2017, McIlroy et al., 2013). DNT-1, DNT-2 and DNT-3 are cleaved intracellularly by furins, whereas Spz-1 is cleaved extracellularly by a Ser-protease cascade that culminates in the activation of Easter or Serine Protease E (SPE), that cleave Spz (Cho et al., 2012, Coutinho-Budd et al., 2017, Foldi et al., 2017, Jang et al., 2006). Importantly, over-expression of cleaved, mature forms of Spz-1, DNT-1 and DNT-2 are functional in vivo, most likely acting cell-autonomously (Hu et al., 2004, Sun et al., 2024, Ulian-Benitez et al., 2017, Zhu et al., 2008). Over-expressed full-length forms of DNT-2 and DNT-3 mimic the natural context, result in natural cleavage and secretion, and function non-autonomously and autocrinely as secreted ligands (Coutinho-Budd et al., 2017, McIlroy et al., 2013, Sun et al., 2024, Ulian-Benitez et al., 2017, Zhu et al., 2008). When cleaved, DNT-1 and DNT-2 promote cell survival (this work, and (Sun et al., 2024, Zhu et al., 2008). In full-length form, DNT-1 can promote JNK signalling, conducive to cell death, whereas DNT-2FL is naturally cleaved to result in mature DNT-2 that promotes cell survival in all contexts tested, including embryos, pupae (this work) and adult brains (Foldi et al., 2017, McIlroy et al., 2013, Sun et al., 2024, Zhu et al., 2008).

Our data indicate that DNT-2 is a neurotrophic factor produced by Mi1 or related medulla neurons, that binds the Toll-2 receptor in L1 lamina neurons, to maintain cell survival, enable appropriate targeting at M1 medulla layer and regulate L1 dendritic complexity. This strongly suggests that lamina neurons receive DNT-2 ligand as they reach Mi1 medulla neurons, that DNT-2 is taken up retrogradely by the connecting L1 neurons, and that establishing connectivity maintains their survival and helps refine their axonal and dendritic projections. Our findings are consistent with prior reports that had shown the maintenance of cell survival to be required during neural circuit formation. The ligand Jeb is produced by R1-R6 photoreceptor axons to activate the receptor Alk located in L3 lamina neuron dendrites, and control their survival during 20-40h APF, at the peak of cell death (Pecot et al., 2014, Togane et al., 2012). Upon loss of function for of Jeb or Alk, L3 neurons are eliminated by apoptosis as their survival could be rescued by over-expressing the inhibitor pf apoptosis p35 (Pecot et al., 2014). L5 neuronal survival is also maintained non-autonomously: two L5 neurons are initially produced, but only the one that receives Col4A1 and EGF signalling activates MAPK and is maintained alive and differentiates into L5 neurons (Fernandes et al., 2017, Prasad et al., 2022). In the medulla, Dm8 medulla neurons are produced in excess and are eliminated during connectivity to their R7 inputs (Courgeon and Desplan, 2019a). This is enabled by the cell surface molecular tags DIPγ in yDm8 binding Dpr11 in yR7, during synaptic matching (Courgeon and Desplan, 2019a). Furthermore, survival of Dm12 and Dm14 neurons depends on interactions between DIP-α and its binding partners, Dpr10 and Dpr6 (Xu et al., 2022, Xu et al., 2018). Loss of interactions between the DIP-α and Dpr10 and Dpr6 causes death of Dm4 and Dm12 neurons and mistargeting of Dm12 from M3 layer to M8 (Xu et al., 2018). Importantly, the maintenance of cell survival takes place during connectivity, and enables synaptic matching between connecting neurons. By contrast, it has also been proposed that apoptosis plays a minor role in cell number control during visual system development, depending instead on cell proliferation and spatial patterning through Dpp/BMP signalling (Malin et al., 2024). However, those findings were based on events taking place at the larval third instar wandering stage, when proliferation and spatial patterning are prevalent, whereas apoptosis peaks in pupa. Altogether, our findings show that the non-autonomous maintenance of neuronal survival is required for appropriate visual system development.

Conceivably, the functions of DNT-2 in promoting neuronal survival and modifying connectivity patterns could reflect independent functions. In fact, over-expression of *DNT-2* both in the adult brain and at the neuromuscular junction, long after circuitry is established, can increase arborisations and induce synaptogenesis (Sun et al., 2024, Ulian-Benitez et al., 2017). Therefore, DNT-2 can induce changes in neuronal shape and connectivity also independently of regulating cell survival. This is also the case for mammalian neurotrophins, which promote axonal and dendritic growth, and synaptogenesis, as well as cell survival (Lu et al., 2005, Park and Poo, 2013). Our data here would suggest that DNT-2 and Toll-2 are not required for axon guidance, as surviving L1 neurons in *DNT-2* or *Toll-2* loss of function genotypes can reach their targets. However, *Toll-2* mutant MARCM clones generated in the pupa result in a dramatic loss of lamina neuron dendrites and aberrant axonal navigation in the medulla, as well as widespread neuronal loss (Li et al., 2020). Here, we have shown that altering DNT-2 and Toll-2 levels impairs appropriate targeting of surviving L1 neurons at the M1 layer – where L1 neurons connect Mi1 neurons - and modifies L1 dendritic complexity. Importantly, connectivity between L1 and medulla neurons takes place between 20-48h APF, during the period of naturally occurring cell death, and spontaneous activity in the optic lobe takes place at 48h, meaning at least some circuits are connected by then (Kurmangaliyev et al., 2020). These data indicate that DNT-2 is required to maintain connecting neurons alive and enables the stabilisation and refinement of their connections.

When altering DNT-2 or Toll-2 levels, L1 axonal terminals in the medulla were misrouted, rather than being confined to a single column. This is reminiscent of the phenotypes caused by alterations in Dscam and Fez levels (Millard et al., 2007, Peng et al., 2018). In the Drosophila visual system, R7, R8, and

L1-L5 neurons form connections in a single column within the medulla’s layers, each containing one axon of each of these cells (Neriec and Desplan, 2016). The cell adhesion molecule Dscam2 drives repellent interactions between L1 neurons, restricting them to a single column (Millard et al., 2007). Any alteration in Dscam2 levels resulted in misrouting of L1 axonal terminals into nearby medulla columns (Millard et al., 2007). Similarly, both over-expression for *DNT-2* and *Toll-2* knock-down resulted in aberrant targeting at the M1 layer. By contrast, this was likely caused by inappropriate levels of DNT-2 depleting the information content normally carried out by secreted ligands in limiting amounts.

Neurotrophins are also required to maintain neuronal survival and enable connectivity in the vertebrate visual system. For example, BDNF with its receptor TrkB promote neuronal survival (Cohen-Cory and Fraser, 1994), as well as axonal arborization and synapse maturation in retinal ganglion cells, during retinotectal synaptic connectivity (Marshak et al., 2007, Sanchez et al., 2006). Disrupting TrkB signaling altered the branching and synaptic maturation of presynaptic axon arbors and suggested that presynaptic TrkB signaling in retinal ganglion cells is a key determinant in the establishment of visual connectivity (Marshak et al., 2007). The evolutionarily conserved functions of DNT-2, and to some extent of DNT-3, during visual system development supports the notion that neurotrophism is a fundamental property of nervous system formation, operating in Drosophila as well as in vertebrates.

To conclude, our findings show that during the apoptosis wave sweeping across the visual system in pupal development, survival of Toll-2+ lamina L1 neurons depends on DNT-2. They also suggest that the survival of Toll-2+ medulla neurons that project within the medulla and to the lobula complex also depends on DNT-2. The findings also suggest that DNT-2 could also influence neurons expressing its canonical receptor Toll-6. Furthermore, DNT-3 also promotes cell survival in the optic lobe and it might interact with Toll-8 or other Tolls to regulate the survival of other lamina and medulla neurons in the optic lobe. In this way, different lamina neurons expressing different Tolls or combinations of Tolls, could be regulated by DNT-2 or DNT-3 or both. As DNT-2 is secreted in medulla neurons and Toll-2 is expressed along neurons that connect in the medulla (e.g. L1, Mi1, Tm3, Dm9, T4), DNT-2 could help keep connecting neurons together during dynamic cellular events in development. Perhaps DNT-3 could also operate similarly in an alternative pathway (e.g. L2). Altogether, the regulation of cell survival by DNTs and Toll receptors provides an additional strategy contributing to the robustness of neural circuit development.

## MATERIALS AND METHODS

### Genetics

Please see S1 Table for the list of the stocks used and Table S3 for full genotypes for each experiment. Mutants: *DNT-2*^*37*^ and *DNT-2*^*18*^ are protein null alleles (Foldi et al., 2017, Ulian-Benitez et al., 2017, Sun et al., 2024). *spz3*^*46*^ and *spz3*^*27*^ are loss of function alleles, generated from P-element excision of from *spz-3*^*EY06670*^.

#### GAL4 driver lines

In-frame T2A-fusions generate a shared transcript between the gene of interest and GAL4, that liberate a functional GAL4 protein that reproduces the endogenous expression of the gene of interest. Spliced-in fusion transcript was verified by PCR. Therefore, these drivers reproduce the endogenous expression of the gene of interest. *spz-1*^*MIO2318*^*-T2A-Gal4* was generated by Recombination Mediated Cassette Exchange (RMCE) by inserting T2A-Gal4 in-frame into MIMIC allele *spz-1*^*MIO2318*^. *spz-3/DNT-3-T2A-GAL4* was generated by CRISPR/enhanced homologous recombination to insert T2A-Gal4 in frame within the first intron of the gene (see below). *Spz-4-T2A-GAL4* was generated by RMCE, by inserting T2A-Gal4 in-frame into intronic *spz-4*^*MI15678*^ MIMIC allele. *DNT-2-T2A-Gal4* is a CRISPR/Cas9-knock-in allele, with GAL4 at the start of the gene (Sun et al., 2024). Toll-1-T2A-Gal4 is a CRISP/Cas9 knock-in at the start of the gene (Singh et al., 2025). *Toll-2*^*PTV*^*GAL4* (Li et al., 2020) has a fully functional GAL4 coding region inserted into the attP site of the pTV cassette (Baena-Lopez et al., 2013) that replaces the coding region for Toll-2 (Li et al., 2020). Toll-6GAL4^MIO2127^ was generated by RMCE by inserting GAL4 into the *Toll-6*^*MIO2127*^ MIMIC in the only exon of this intron-less locus (Li et al., 2020). *Toll-8GAL4*^*MD806*^ is a P-element insert-180bp upstream of the Toll-8 start codon (Li et al., 2020). *Nsyb-Gal4* drives GAL4 in all neurons (BDSC#3917). L1-split GAL4: *w; 84A08-p65ADZp attp40; 66A01-ZpGdbd attp2* (gift from the Reiser Lab) (Tuthill et al., 2013).

#### Reporter lines

*UAS-mCD8::GFP*, for membrane tethered *GFP; UAS-histone-YFP*, for YFP-tagged nuclear histone; *10XUAS-myr::GFPattP40* on 2^nd^ (BDSC#32198), *10XUAS-myr::GFPattP2* on 3^rd^ (BDSC# 32197), and 10XUAS-myr::GFP, su(Hw) attP8 on X (BDSC# 32196) whereby a myristoylation tail at the N-terminus of GFP associates it to the plasma membrane; 20XUAS-6XmCherry-HA attp2 on 3^rd^ (from BDSC#52268) for cytoplasmatic expression of a hexameric form of mCherry.

#### UAS for gene over-expression, knock-down and epistasis

*UAS-DNT-2FL* (full length) and *UAS-DNT-2CK* (signal peptide plus cystine-knot, mature form) (Foldi et al., 2017, Sun et al., 2024, Ulian-Benitez et al., 2017, Zhu, 2013); *UAS-spz3FL* (full length) and UAS-spz3CK (signal peptide plus cystine-knot, mature form)(this work); *UAS Toll-2 RNAi*^*v36305*^ (VDRC36305) (Li et al., 2020). To select pupae of the desired genotype, the fusion balancer SM6aTM6B marked with Tb- and which segregates the second and third chromosomes together, was used. All experiments were carried out at 25°C unless otherwise indicated.

#### Multi Colour Flip Out (MCFO) clones

*hs-FLPG5.PEST;;10xUAS (FRT. stop) myr::smGdP-OLLAS 10xUAS (FRT. stop) myr::smGdP-HA 10xUAS (FRT. stop) myr::smGdP-V5-THS-10xUAS (FRT. stop) myr::smGdP-FLAG* (BSC64086) flies were crossed to *w; Toll-8GAL4, w;Toll-6*^*MIO2127*^*GAL4* or *w; Toll-2*^*pTV*^*GAL4* flies and bred at 25°C. L3 wandering larvae where watched until pupariation, 20 white pupae were fished out at once and moved to a new vial, placed at 25° for 24h (24h after puparium formation, APF), heat shocked in a water bath at 37° for 5 minutes, then placed at 25° until they reached 72h APF, then dissected, fixed and stained.

### Molecular biology

*Spz-3/DNT-3-T2A-GAL4* was generated by CRISPR/enhanced homologous recombination, by inserting T2A-GAL4 from plasmid pT-GEM(1) phase 1 (AddGene 62893) into the first intron of the *spz-3/DNT-3* coding region (between the first and second coding exons), to create a spliced in-frame fusion transcript with the 5’-terminus of *spz-3/DNT-3* mRNA and bearing T2A-GAL4 mRNA in frame. The guide RNA targeted the same intron (gRNA primers: GTCGTTTGGGTCGCTCGATGTCT and AAACAGACATCGAGCGACCCAAAC, Table S2), and it was designed using the Optimal Target Finder. BbsI enzyme sites were added to the oligos used to generate the gRNA. The gRNA was cloned into pU6.3 using conventional ligation. 1kb of genomic DNA were amplified for each homology arm, from nos-Cas9 genomic DNA using Q5 enzyme (Promega), using primers: 5’ homology arm: GGTATACCGGTCGAATAAGTGACTCAAGCAGAC and ATAGCGGCCGCAGTATCTGAGTTTTGGTCTTG; 3’ homology arm: TATGGTACCTCTTGGCGCGGCACTCAAGT and CACACTAGTCTCATGCCGGCGAACCTATC. To clone the homology arms into the pT-GEM(1) plasmid, AgeI and NotI cut sites were added at the extreme of the 5′homolgy arm, and KpnI and SpeI cut sites were added to 3′ homology arm ends. Both constructs were injected in flies expressing Cas9. After the selection of transformants carrying the 3xP3-RFP marker, stocks were balanced and the red fluorescent marker removed by CRE-recombinase. Cleaved UAS-spz-3CK/DNT-3CK was made by cloning the signal peptide of spz-5 (100bp) inserted between EcoRI and BglII, followed by the cystine-knot domain of spz-3 (primers: spz-5 SP: CGGAATTCATGCAAATCGACGGCGAATGA with EcoRI site and GAAGATCTCGAGCTGTGGGCGGCTACTGT with BglII; start of spz-3 Cysknot: GAAGATCTGCCGGAGGAAGTCGAAATAGA with BglII site and end CCGCTCGAGCAGAGTCAGGTAATCTAGGGA with XhoI), into pUAS-attB, and Phi-C31 transgenesis into 86Fa. Full length UAS-spz-3FL/DNT-3-FL was generated by Gateway cloning, from cDNA full length clone RE22741. Using primers GGGGACAAGTTTGTACAAAAAAGCAGGCTCGCTAGCATATTTCGCACGCCC and GGGGACCACTTTGTACAAGAAAGCTGGGTCGGGATTACATCTACAGACAC (Table S2), the coding sequence of spz-3/DNT-3 was amplified and the PCR product was inserted with a BP reaction into attB sites of the pDONR221 vector to generate an entry clone with attL sites. Next, *spz-3/DNT-3-FL* was inserted into attR sites of destination vector *pUAS-GW-attB* with an LR reaction, to generate the final expression clone with attB sites and the final construct was inserted with Phi-C31 transgenesis into attP2 landing site. 3xP3-RFP was removed with CRE-recombinase.

#### Analysis of scRNAseq from published databases

We analysed single-cell RNA sequencing data from the study by (Kurmangaliyev et al., 2020) (GSE156455), specifically the pre-processed dataset covering 24–96 hours after puparium formation (APF), accessed via Zenodo (https://doi.org/10.5281/zenodo.4264808). All downstream analyses were performed using Seurat v3 in R. The data were imported as a Seurat object, and cells corresponding to specific timepoints (e.g., 24 h, 36 h) were subsetted based on the provided metadata. Dimensionality reduction was carried out using principal component analysis (PCA), followed by Uniform Manifold Approximation and Projection (UMAP) embedding computed on the first 30 principal components. Cluster annotations provided by the original authors were used for all cluster-level analyses and visualisations. For feature visualisation, FeaturePlot and custom ggplot2-based UMAP plots were generated, incorporating manually defined colour palettes and cluster-specific labelling where appropriate.

### Immunostaining

Immunostainings in the pupal brains were carried out following standard protocols. Pupae were staged by picking the white pupa at the end of L3-wandering. For each stage (24h, 48h, 72h APF) the date and time of fishing were recorded to start dissection at the appropriate timepoint. Dissections were carried out in PBS with forceps in 20 minutes windows. Dissected brains were placed in 4% formaldehyde and kept on ice: for Dcp-1, brains were fixed for 20 minutes; for all other stainings brains were fixed for 35 minutes. After fixation, brains were blocked for 2 hours in normal goat serum and primary antibodies were incubated overnight. This was followed by incubating in secondary antibodies and washes. Primary antibodies used were: mouse anti-24B10 (DSHB) at 1:250 dilution; rabbit anti-GFP (Thermo Fisher) at 1:250; rabbit anti-Dcp-1 (Cell Signalling) at 1:250; rat anti-N-cadherin (MAb DN-Ex) at 1:250; chicken ant-HA (Aves) at 1:100. Secondary antibodies used: Alexa Donkey-anti-Rabbit 488 at 1:250; Alexa Goat-anti-mouse 647 1:250; Alexa Goat-anti-Rat 647 1:250; Alexa Goat-anti-Chicken 647 1:250. Details of primary and secondary antibodies used, their origin and working dilutions are also provided in Table S3.

### Microscopy and imaging

#### Mounting of optic lobes

for visualisation of expression profiles (Figures 1,2) optic lobes were mounted laterally; for all other data they were mounted horizontally.

#### Laser scanning confocal microscopy was carried out using

Figure 1, 2: Zeiss (LSM 710) at resolution of 512×512 or 1024×1024 pixels, speed 6 or 7, 20× objective and step 1μm. For DCP-1 stainings in Figure 3, Figure 4A,B and Figure S9, samples were scanned using Zeiss LSM900 with Airyscan 2, at 512×512 pixels, using a 25x oil objective with 0.9x zoom, speed 7 and step 1μm. For Histone-YFP in Figure 4C-F, Zeiss LSM900 was used at 1024×1024 pixels, 25x oil objective 0.8x zoom, speed 7 and step 1μm. Figures 5 and 6 data were acquired with Zeiss LSM900, at 1024×1024, no zoom and 2x averaging for axons (Figure 5) and 2x zoom and 8x averaging for dendrites (Figure 6), step 1μm.

#### Quantification of apoptosis

Anti-Dcp1 labelled pupal cells do not have a regular morphology and are not suitable for automatic cell counting with DeadEasy nor Imaris Spot Function, thus volume intensity measurements were made instead. To analyse Dcp1 signal volume intensity using Imaris (Figure 3, Figure S9 and Figure 4A,B) a 3D-ROI was selected for either the lamina or optic lobe minus lamina, followed by creating a surface to mask the ROI. Next, the Wizard Wand was selected to render the volume ROI, setting a threshold to match the DCP-1 signal whilst eliminating small non-specific background dots. Care was taken to ensure that samples were of equivalent quality prior to processing and that the thresholding worked consistently across samples. Once adjusted, the Split Touching Objects feature was enabled to isolate DCP-1 signal from background. ROI volume signal intensity was selected.

#### Automatic cell counting

Nuclei labelled with histone-YFP did not require antibody stainings. 3D-Regions of Interest (ROIs) through the stack of confocal images covering the entire optic lobe of each specimen were selected using Imaris. The ROI.tif files were then opened in ImageJ and automatic cell counting was carried out using DeadEasy Optic Lobe, as previously described (Li et al., 2020), and publicly available at UBIRA: https://edata.bham.ac.uk/1213/.

#### Quantification of signal intensity using Fiji (Figure 4E,F)

maximum projections of optical sections through the anterior (corresponding approximately to the medulla) or posterior (corresponding approximately to the lobula complex) optic lobes mounted horizontally were generated. An ROI was traced around the optic lobe excluding the lamina, and signal intensity (Integrated Density, IntDen) was measured using the “Measure” tool in Fiji. ROI IntDen was normalized by subtracting the background IntDen, obtained from the mean intensity density of a small ROI within the tissue but not containing hisYFP+ cells multiplied by the ROI area.

#### Lamina L1 axonal misrouting

Lamina L1 neurons were visualized with *L1-splitGAL4>GFP* and dissected pupal CNSs were counterstained with 24B10 to visualize all photoreceptor axons. Using Fiji, Z-projections of 1-5 optical 1μm sections were generated to trace individual L1 projections. 24B10+ photoreceptors R7 and R8 project into the medulla arranged in neat columns that do not intersect with each other. L1 neurons target in medulla layers M1 and M5. If the axon of an L1 neuron was constrained within a single column, this was considered “a wild-type phenotype”. If the axon crossed into neighbouring columns on either side, this was considered “a misrouting phenotype”, as in (Millard et al., 2007). Scoring was carried out manually.

#### Quantification of L1 dendrite volume

Amira was used to measure the volume of the L1 neuron dendrites in 3D. Stacks of confocal images of L1 neurons were imported into Amira. Within the segmentation section, the magic wand was used to select the dendritic spines, and thresholding was applied to faithfully trace them, across the stack. Following the selection, volume retendering was applied to select and measure the full dendrite volume. Dendrite volume signal intensity was selected.

### Statistical analysis

Statistical analysis was carried out using GraphPad Prism. The confidence interval was 95%, and significance set at p<0.05. For quantitative continuous data, D’Agostino or Shapiro-Wilk normality tests were carried out. Data that were normally distributed were analysed using Student t-test for comparisons between two genotypes. When more than two samples were being compared, equality of variances was tested (Levene’s or Barlett’s test), and if equal, One-Way ANOVA was used, followed by a post-doc Dunnett test for multiple comparisons to a fixed control. When normally distributed data did not pass the equality of variance test (Brown-Forsythe), Welch ANOVA weas used, followed by post-doc Dunnett test for multiple comparisons to a fixed control. For quantitative data that were not normally distributed, Kruskal-Wallis was used for comparisons between more than two genotypes, followed by post-hoc Dunn’s test for multiple comparisons to a fixed control. Categorical data were analysed with Fisher’s exact tests and post-hoc Bonferroni multiple comparisons corrections. For further details, p values, genotypes and sample sizes, please see Table S4.

## ACKNOWLEDGEMENTS

We thank Chris Bunce, Yun Fan, Filipe Pinto-Teixeira, Natalia Sanchez-Soriano and members of our lab for discussions and feedback; J.C. Tuthill and M.B. Reiser for the kind gift of fly stocks; Bloomington Drosophila Stock Centre for stocks; DSHB (Iowa) for antibodies; AddGene for plasmids; FlyBase for facilitating our research. This work was funded by a PhD Scholarship by the Ministry of Higher Education and Scientific Research, UAE. to N.A., and BBSRC Project Grant BB/R017034/1, MRC Career Establishment Grant, Wellcome Trust Project Grant 088583/Z/09/Z and Wellcome Trust Investigator Award 223197/Z/21/Z to A.H.

## AUTHOR CONTRIBUTIONS

N.A., F.R.C, B.Z., S.A-F, M.M, G.L., M.M., and A.H. designed and/or executed experiments; N.A., F.R.C, B.Z., S.A-F, M.M., A.L. and A.H. analysed data; N.A. and A.H. wrote the manuscript, and all authors provided feedback. A.H. conceived and directed the project.

## FIGURE LEGENDS

**Supplementary Figures S1 to S6 Combined UMAPs** showing the expression of *spz-1, -3 (DNT-3), -4* and *-5 (DNT-2)* and *Toll-1, -2 (18w), -6* and *-8 (Tollo)* in distinct cells over time.

**Supplementary Figures S7 scRNAseq: expression levels of DNT-2 (spz-5) and 18w (Toll-2) over time in each cell type (Kurmangaliyev et al dataset)**. Arrows indicate L1 and L3 lamina neurons, some medulla neurons (Mi1, Mi4 and Mi9) and T4/T5.

**Supplementary Figure S8 DNT-2 and DNT-3 can maintain cell survival during optic lobe development also outside the lamina. (A**,**B)** Over-expression of *spz-3/DNT-3* in Toll-8 neurons *(Toll-8>spz-3/DNT-3)* and **(C-D)** of *DNT-2/spz-5* in all neurons *(nsyb>DNT-2)* can rescue naturally occurring cell death outside the lamina, in optic lobe regions comprising the medulla and lobula complex. Apoptotic cells labelled with anti-Dcp1 at 24h APF. Images in **(A**,**C)** are projections of the anterior optic lobe corresponding to the medulla and the posterior optic lobe corresponding to the lobula complex, to show Dcp1+ apoptotic cells. Dotted lines in **(A**,**C)** indicate indicate ROI excluding the lamina used for measuring signal volume in 3D using Imaris. Unpaired Student t tests: **(B)** ***p<0.0001; **(D)** *p<0.05 For further details, see Table S4. Scale bar: 50μm.

**Supplementary Table S1 Key resources: List of Drosophila fly stocks used**

**Supplementary Table S2 Key resources: List of primers used**

**Supplementary Table S3 Key resources: List of primers antibodies**

**Supplementary Table S4 Statistical analysis, sample sizes and genotypes**.

