## Supplementary material for "The neurotrophin DNT-2 regulates cell survival and connectivity via the Toll-2 receptor during visual system development of *Drosophila*": Table S1

**Table S1	Stock list**

| **What?** | **Genotype** | **Source** |
| --- | --- | --- |
| Oregon | Oregon R |  |
| nsybGAL4 on 3^rd^ | w; nsybGAL4 attp2 | BDSC#39171 |
| 10XUASmyrGFP on 2^nd^ | P{10XUAS-IVS-myr::GFP}attP40 | BDSC# 32198 |
| 10XUASmyrGFP on 3^rd^ | P{10XUAS-IVS-myr::GFP}attP2 | BDSC# 32197 |
| 10XUASmyrGFP on X | P{10XUAS-IVS-myr::GFP}su(Hw)attP8 | BDSC# 32196 |
| UASHistoneYFP on 2^nd^ | w; UAS histone YFP (2) |  |
| MCFO on 3^rd^ | w hsFLP:PEST ;; HA-V5-FLAG-OLLAS MCFO | BSC #64086 |
| UAS mCD8 GFP on 2^nd^ | w; UAS mCD8 GFP |  |
| Toll-1 Gal4 on 3^rd^ | w;+/(CYO);Toll-1-T2A-Gal4 [CRISPR]/ TM6B | Hidalgo lab  Singh et al 2025 |
| Toll-2 Gal4 on 2^nd^ | yw; Toll-2^pTV-^Gal4/CyO | Hidalgo lab  Li et al 2020 |
| Toll-6 Gal4 on 3^rd^ | yw; Toll-6^MI02127^ Gal4/TM3 | Hidalgo lab  Li et al 2020 |
| Toll-8 Gal4 on 3^rd^ | W; Tollo^MD806^Gal4/ TM6B | BDSC #36548 |
| DNT-2 Gal4 on 3^rd^ | w; DNT-2 [CRISPR] T2A-GAL4/MKRS | Hidalgo lab  Sun et al 2024 |
| spz-3 GAL4 on 2^nd^ | w; spz-3-T2A-GAL4/CyO | This work  Hidalgo lab |
| spz-4 GAL4 on 2^nd^ | yw; spz-4^MI15678^-T2A-Gal4 / SM6a | This work  Hidalgo lab |
| spz-3^27^ LOF mutant | w; spz-3^27^/ CyO | This work  Hidalgo lab |
| spz-3^46^ LOF mutant | w; spz-3^46^/ CyO | This work  Hidalgo lab |
| DNT-2^18^ LOF mutant | DNT-2^18^ | Hidalgo lab  Sun et al 2024 |
| DNT-2^37^ LOF mutant | DNT-2^37^ / TM6B | Hidalgo lab  Sun et al 2024 |
| UAS-spz-3FL on 3^rd^ | w; UAS spz-3FL Full length @attP2 | This work  Hidalgo lab |
| UAS-spz-3CK on 3^rd^ | w; UAS spz-3 Cysknot 22A @86FA | This work  Hidalgo lab |
| UAS-DNT-2FL on 3^rd^ | w; UAS DNT-2 FL 22A @86FA | Hidalgo lab  Zhu et al 2008 |
| UAS-DNT-2CK on 3^rd^ | UAS DNT-2 CK6A @86FA | Hidalgo lab  Zhu et al 2008 |
| UAS-Toll-2RNAi on 2^nd^ | W; UASToll-2RNAi[GD36305]/CyO | VDRC36305 |
| L1-Split-GAL4 | R48A08AD;R66A01DBD | Tuthill Reiser Lab |
