## Supplementary material for "The neurotrophin DNT-2 regulates cell survival and connectivity via the Toll-2 receptor during visual system development of *Drosophila*": Table S2

| **Original name** | **Construct or what for** | **Primer Sequence 5🡪3** |
| --- | --- | --- |
| Sense gRNA | Spz3-T2A-Gal4 | GTCGTTTGGGTCGCTCGATGTCT |
| Antisene gRNA | Spz3-T2A-Gal4 | AAACAGACATCGAGCGACCCAAAC |
| Fw-Spz3 5’HA | Spz3-T2A-Gal4 | GGTATACCGGTCGAATAAGTGACTCAAGCAGAC |
| Rv-Spz3 5’HA | Spz3-T2A-Gal4 | ATAGCGGCCGCAGTATCTGAGTTTTGGTCTT |
| Fw-Spz3 3’HA | Spz3-T2A-Gal4 | tatGGTACCTCTTGGCGCGGCACTCAAGT |
| Rv-Spz3 3’HA | Spz3-T2A-Gal4 | cacACTAGTCTCATGCCGGCGAACCTATC |
| SP-EcoRI-Fw | UAS-Spz3-CK  (SP of spz-5) | CGGAATTCATGCAAATCGACGGCGAATGA |
| SP-BglII-Rev | UAS-Spz3-CK  (SP of spz-5) | GAAGATCTCGAGCTGTGGGCGGCTACTGT |
| Spz-3-CK-BglII | UAS-Spz3-CK | GAAGATCTGCCGGAGGAAGTCGAAATAGA |
| Spz-3-CK-XhoI | UAS-Spz3-CK | CCGCTCGAGCAGAGTCAGGTAATCTAGGGA |
| attB1 F spz-3FL | spz3FL pDONR | GGGGACAAGTTTGTACAAAAAAGCAGGCTCGCTAGCATATTTCGCACGCCC |
| attB2 R spz-3FL | spz3FL pDONR | GGGGACCACTTTGTACAAGAAAGCTGGGTCGGGATTACATCTACAGACAC |
| UAS-Spz3-FLGW (F) New | UAS-Spz3-FL | GAGGGTTCGTACTCCCGTTA |
| UAS-Spz3-FLGW (R) new | UAS-Spz3-FL | CGGCTTCGACGGTCTCCCCG |

**Table S2	List of primers**
