## Supplementary material for "The neurotrophin DNT-2 regulates cell survival and connectivity via the Toll-2 receptor during visual system development of *Drosophila*": Table S3

**Table S3	Key resources: List of antibodies**

| **Antibody** | **Donor** | **Working dilution** |
| --- | --- | --- |
| **Primary** |  |  |
| Anti-GFP | Rabbit | 1:250 |
| Anti-Ncadherin | Rat | 1:250 |
| Anti-24B10 | Mouse | 1:250 |
| Anti-DCP-1 | Rabbit | 1:250 |
| Anti-HA | Chicken | 1:100 |
| **Secondary** |  |  |
| Anti-rabbit 488 | Donkey | 1:250 |
| Anti-rabbit 546 | Goat | 1:250 |
| Anti-mouse 647 | Goat | 1:250 |
| Anti-rat 647 | Goat | 1:250 |
| Anti-Chicken 647 | Goat | 1:250 |
