## Supplementary figures and images for "The neurotrophin DNT-2 regulates cell survival and connectivity via the Toll-2 receptor during visual system development of *Drosophila*"

### Supplementary Figure S1

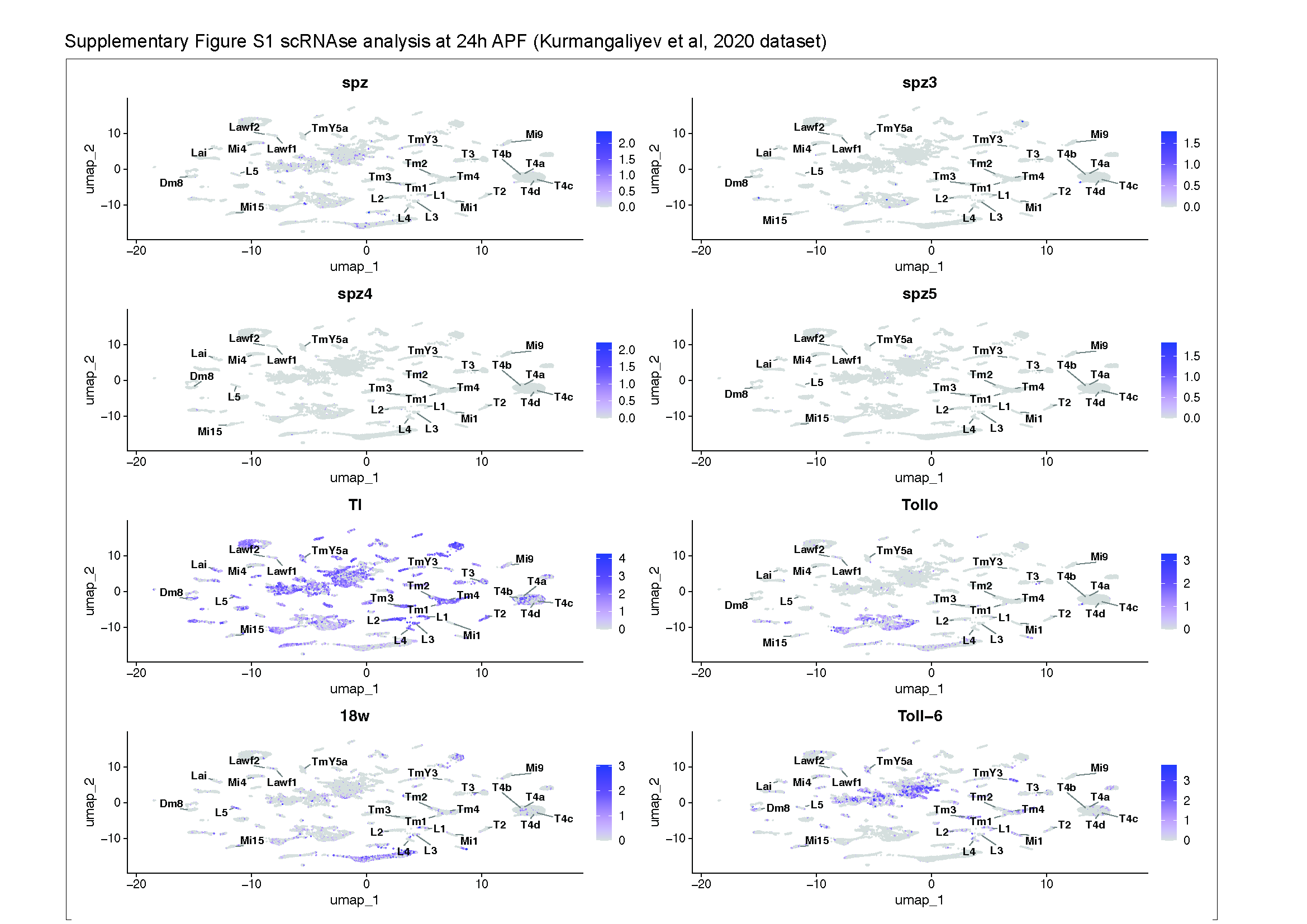

### Supplementary Figure S2

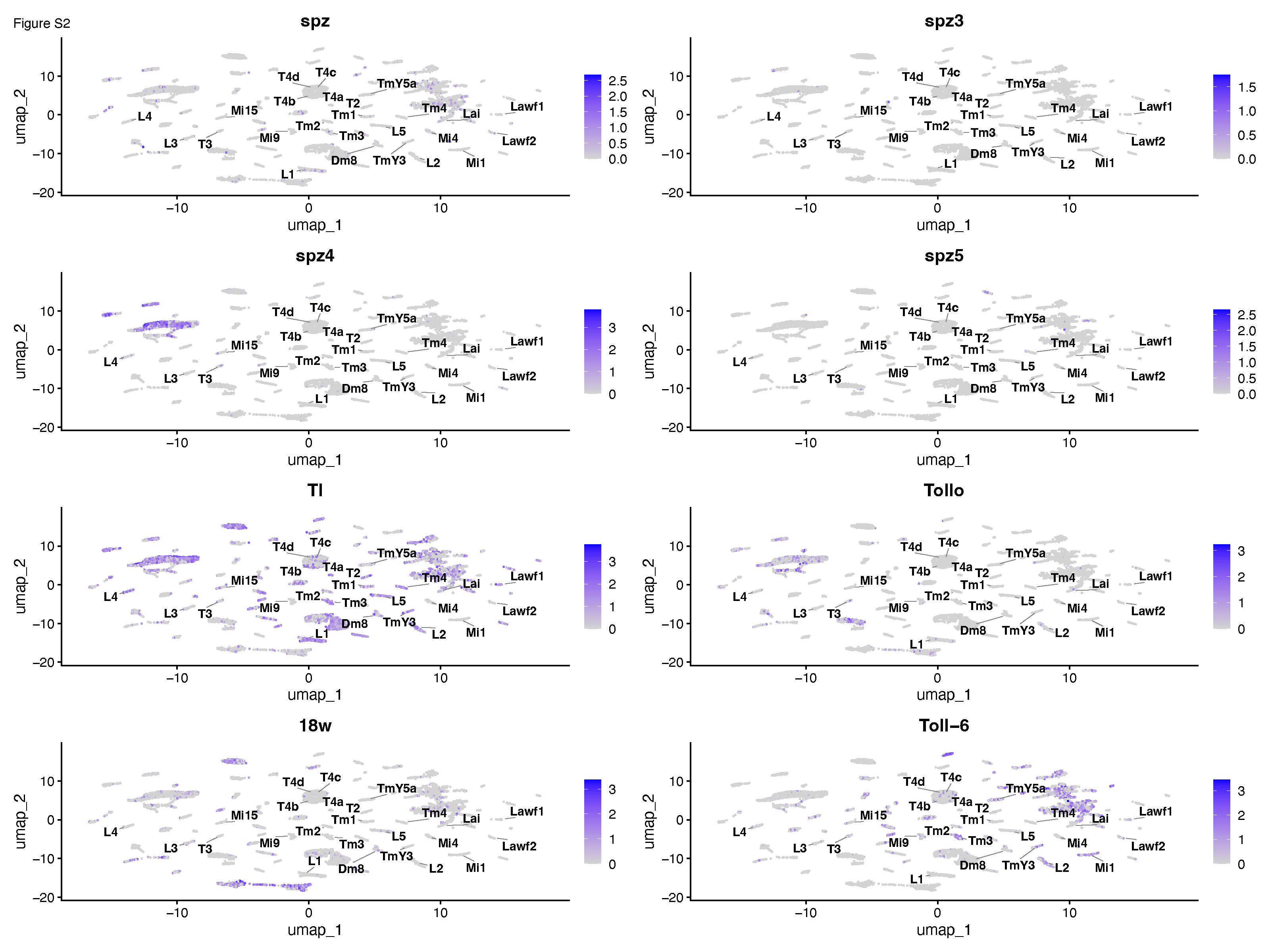

### Supplementary Figure S3

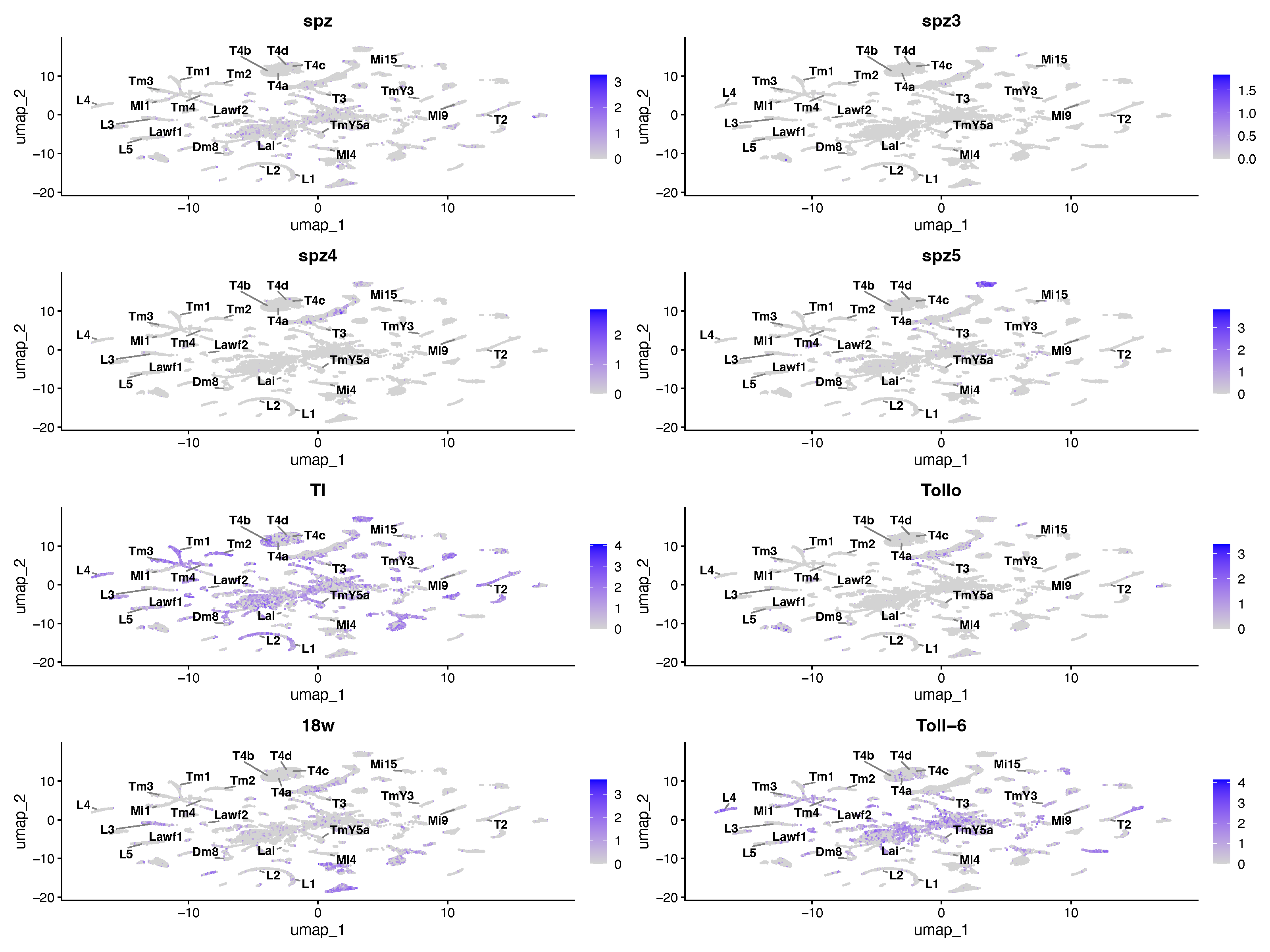

### Supplementary Figure S4

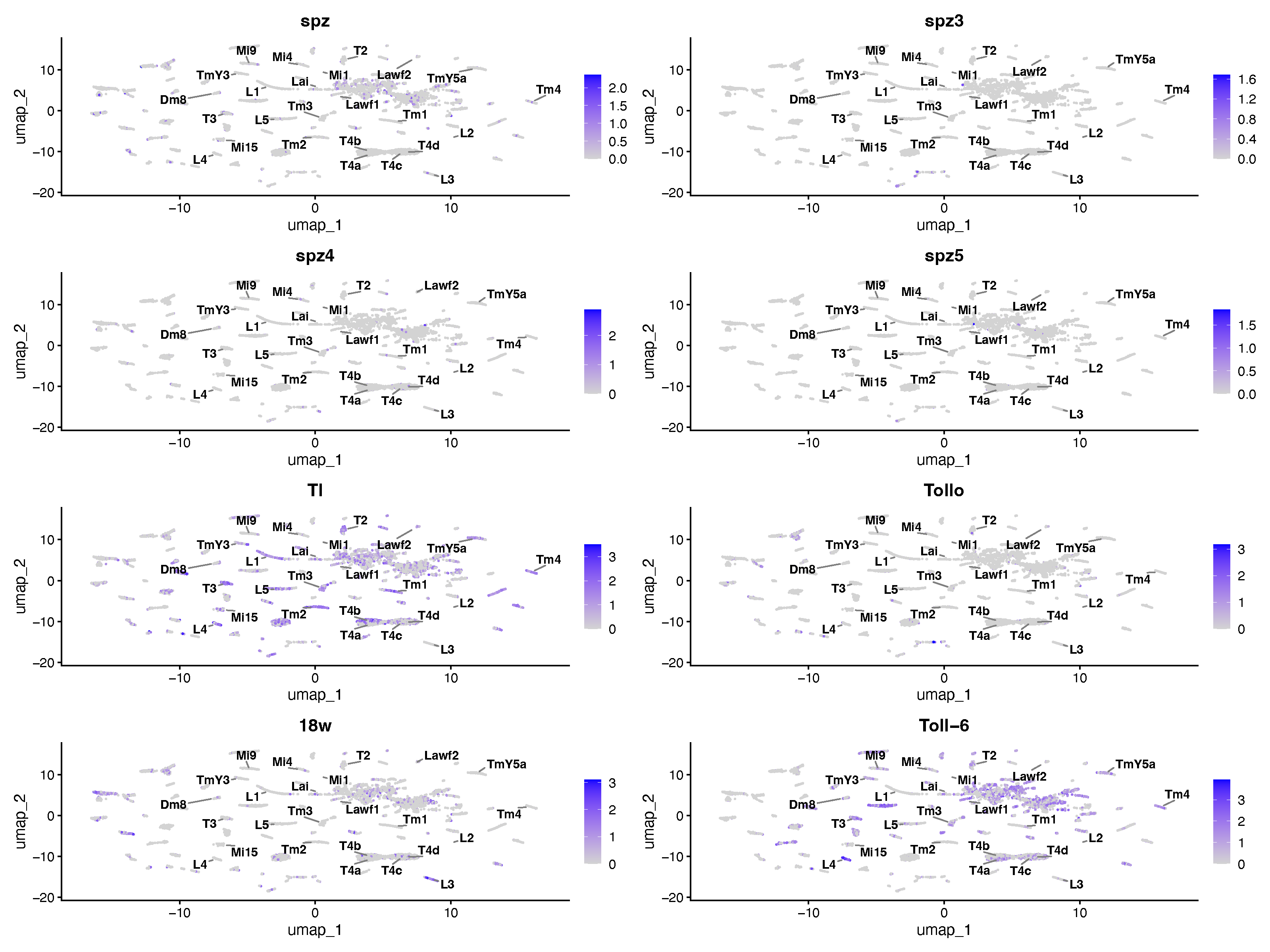

### Supplementary Figure S5

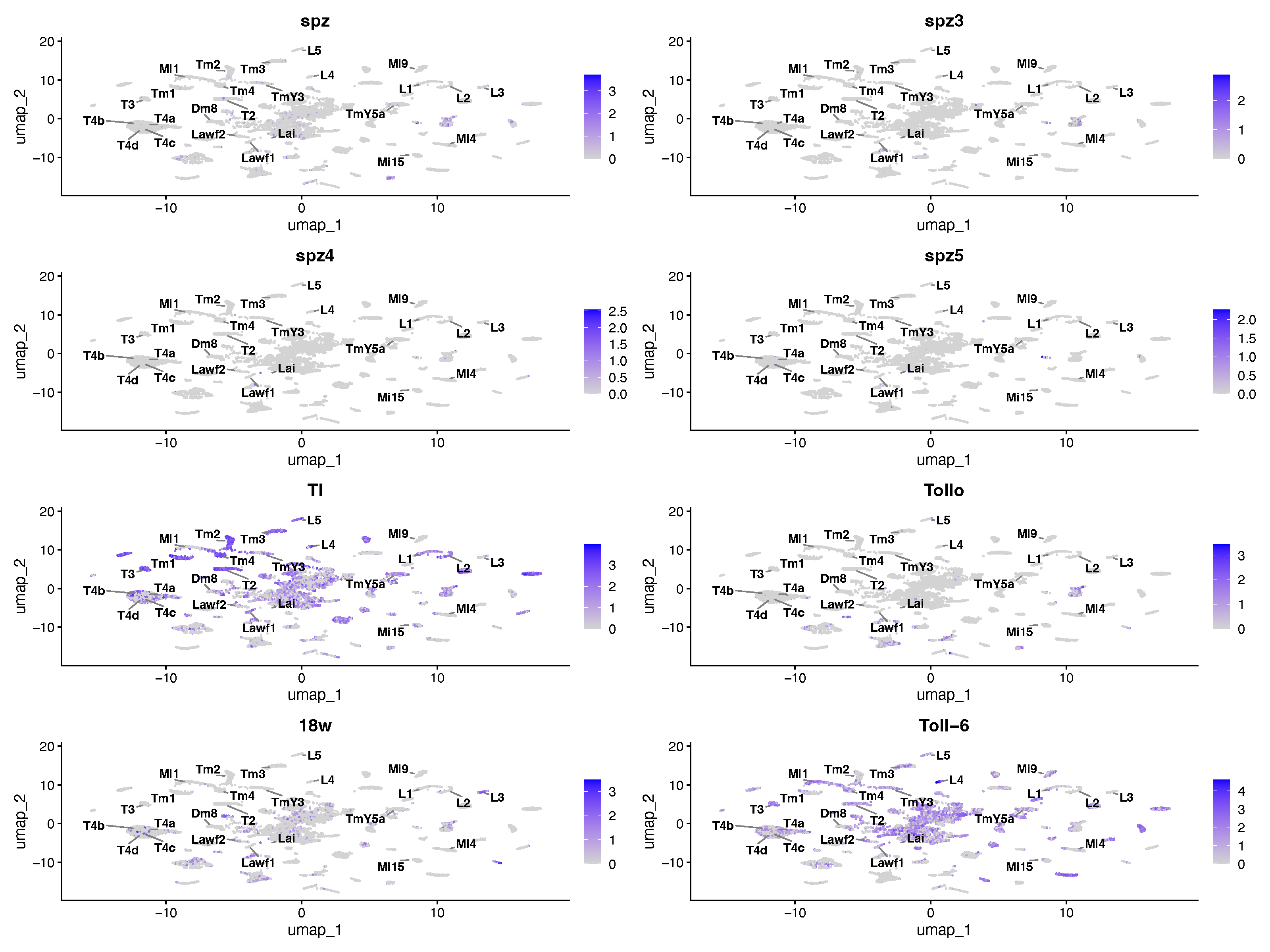

### Supplementary Figure S6

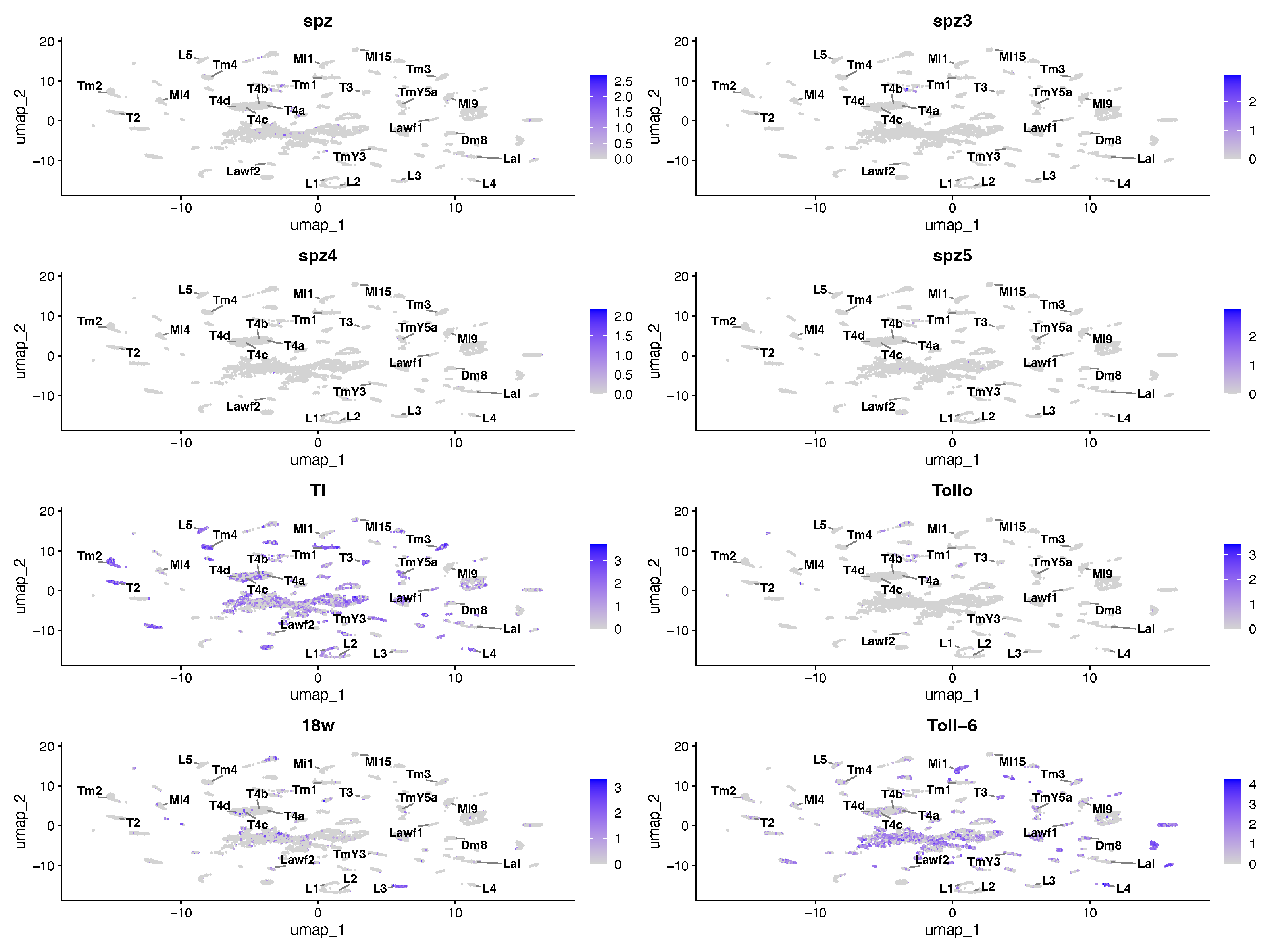

### Supplementary Figure S7

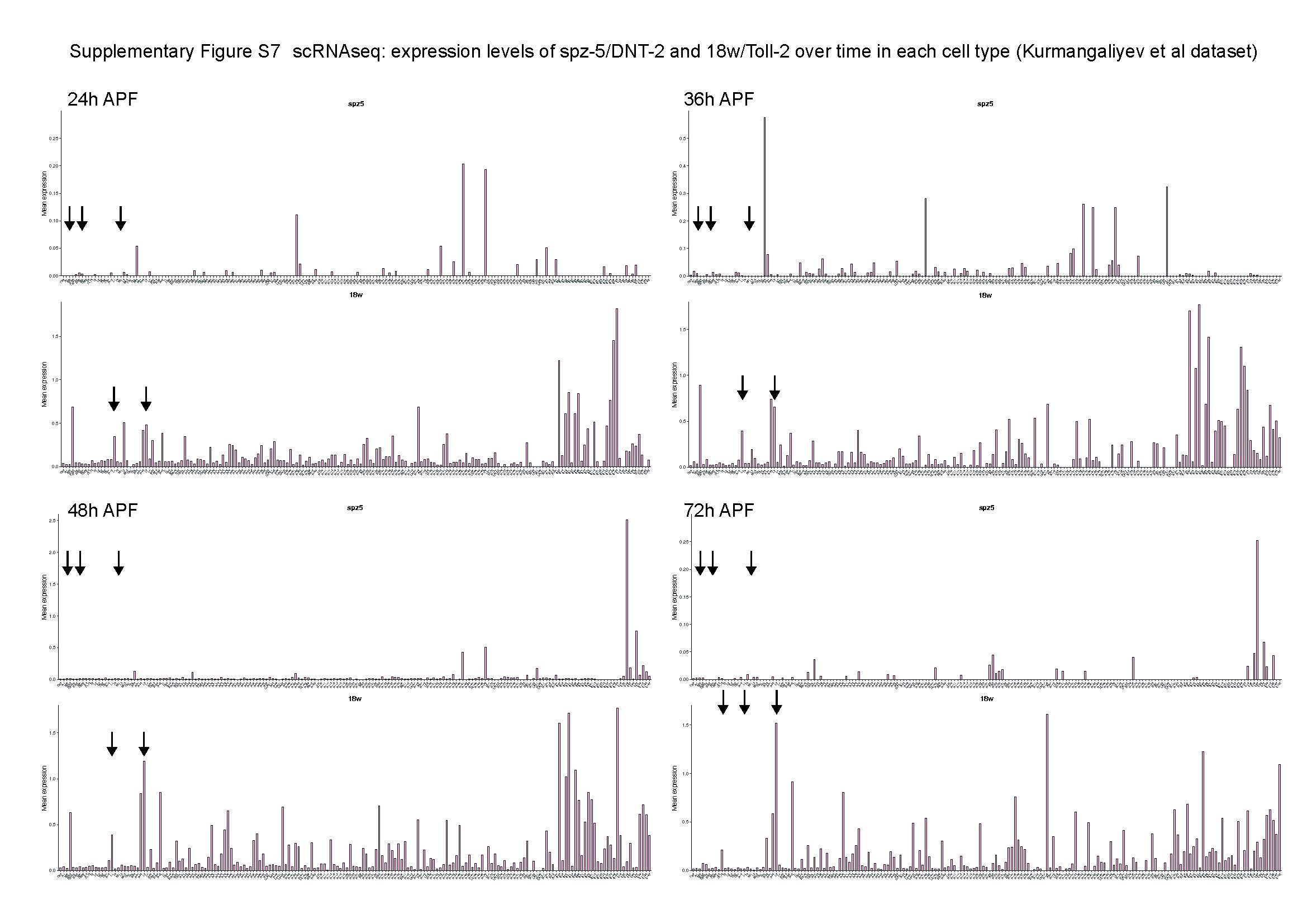

### Supplementary Figure S8

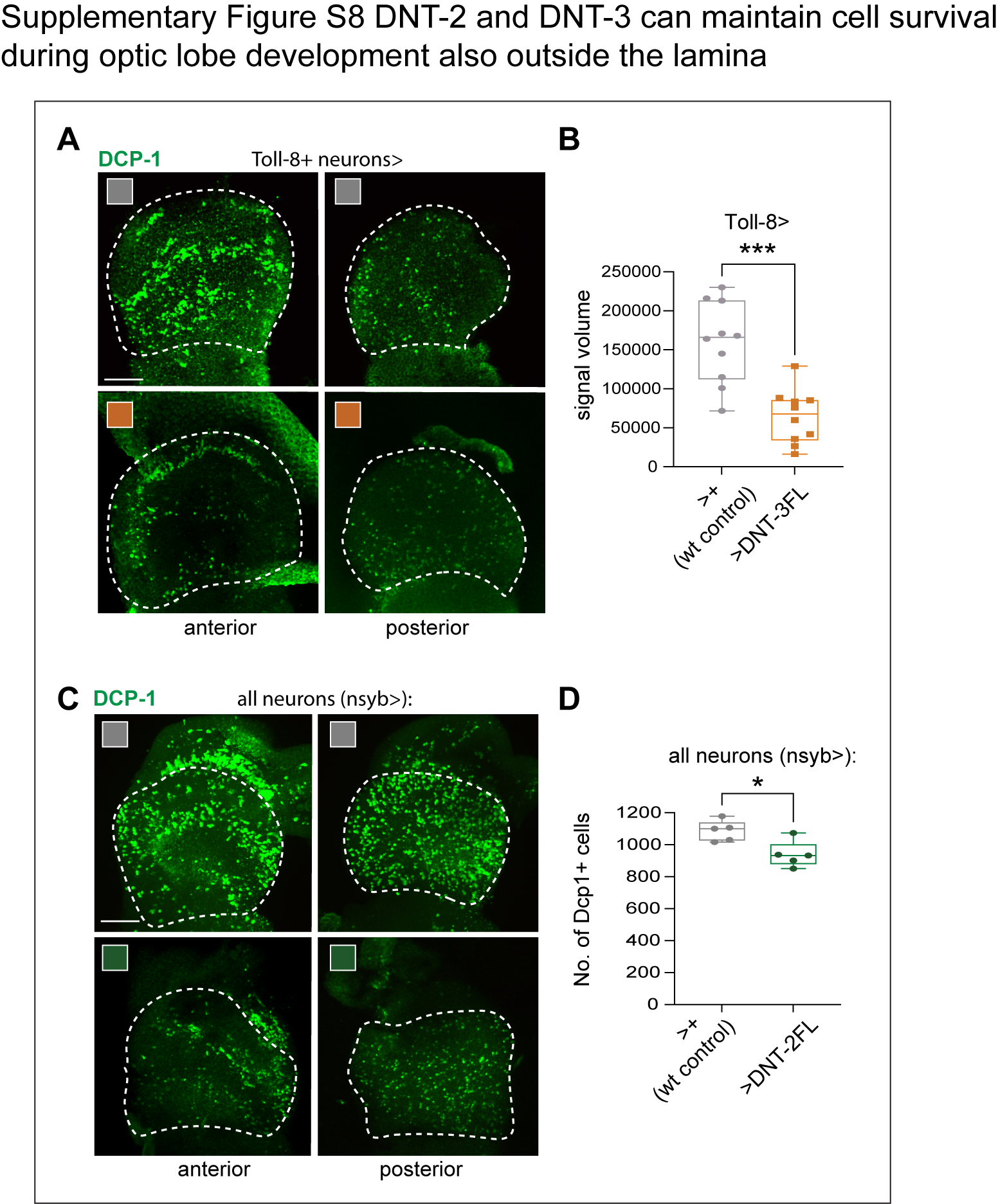
